# Early Hippocampal Sharp-Wave Ripple Deficits Predict Later Learning and Memory Impairments in an Alzheimer’s Disease Mouse Model

**DOI:** 10.1101/596569

**Authors:** Emily A. Jones, Anna K. Gillespie, Seo Yeon Yoon, Loren M. Frank, Yadong Huang

**Author notes:** **Correspondence should be addressed to**: Yadong Huang.

## Abstract

Alzheimer’s disease (AD) is characterized by progressive memory loss, and there is a pressing need to identify early pathophysiological alterations that predict subsequent memory impairment. Hippocampal sharp-wave ripples (SWRs) – electrophysiological signatures of memory reactivation in the hippocampus – are a compelling candidate for doing so. Mouse models of AD show reductions in both SWR abundance and associated slow gamma (SG) power during aging, but these alterations have yet to be directly linked to memory impairments. In aged apolipoprotein E4 knock in (apoE4-KI) mice – a model of the major genetic risk factor for AD – we found that reduced SWR abundance and associated CA3 SG power predicted spatial memory impairments measured 1–2 months later. Importantly, SWR-associated CA3 SG power reduction in young apoE4-KI mice also predicted spatial memory deficits measured 10 months later. These results establish features of SWRs as potential functional biomarkers of memory impairment in AD.

## INTRODUCTION

Alzheimer’s disease (AD) is a form of dementia characterized by progressive cognitive decline that affects 11% of the US population over the age of 65 (Hebert et al., 2013). The continued failure of AD clinical trials has redirected the field towards halting disease progression before symptoms manifest; once memory impairment is detected, it may be too late for treatment to reverse it (Sperling et al., 2011). While there are known genetic and environmental risk factors for AD (Ballard et al., 2011), our ability to predict which individuals will develop the disease, when symptoms will arise, and how rapidly they will progress remains poor. There is therefore a pressing need to identify early pathophysiological alterations which can distinguish later cognitive decline from healthy aging.

To identify early, predictive alterations, we studied a mouse model of ε4 variant of the *APOE* gene, the most common genetic risk factor for AD (Huang and Mucke, 2012; Liu et al., 2013). It has an allelic frequency of 20–25%, yet is found in 65–80% of AD patients (Farrer et al., 1997). The presence of ε4 alleles increases the likelihood of developing AD by age 85 from 10% to 70% in a gene dose-dependent manner (Corder et al., 1993), but does not guarantee AD. Thus, the population of ε4 carriers shows significant individual variation, and predicting that variation could allow preventative treatment to be targeted to the highest risk individuals. Mice with human *APOE* ε4 knocked in at the mouse *APOE* locus (apoE4-KI) have physiologically appropriate patterns and levels of apoE expression (Bien-Ly et al., 2012; Ramaswamy et al., 2005) and recapitulate gender and age effects of apoE4 seen in humans (Andrews-Zwilling et al., 2010; Beydoun et al., 2012; Farrer et al., 1997; Leung et al., 2012). By 16 months of age, female apoE4- KI mice show spatial learning and memory deficits as measured in the Morris water maze (MWM), recapitulating age-dependent memory loss as seen in human ε4 carriers (Caselli et al., 2009) and providing a useful animal model for longitudinal study of memory decline.

A memory-predicting alteration should reflect underlying pathology of AD and be related directly to memory processes. The hippocampus is one of the first sites of AD pathology and is required for the spatial learning and memory processes that falter early in AD (deIpolyi et al., 2007; Mattson, 2004; Morris and Baddeley, 1988; Squire and Wixted, 2011); thus, physiological signatures of hippocampal information processing could provide useful biomarkers. The hippocampal local field potential (LFP) is a particularly appealing candidate. The LFP is an extracellular voltage measurement that reflects the summation of activity patterns from local neurons. The LFP provides real-time measures related to memory processing including consolidation and retrieval (Buzsáki, 2015) and can be repeatedly measured in the same subject; for these reasons, features of the LFP have been previously proposed as potential biomarkers in AD (Goutagny and Krantic, 2013; Palop et al., 2006). We focused on sharp-wave ripples (SWRs), an LFP signature of memory replay. During SWRs, a large population of hippocampal neurons is activated, often in sequences that recapitulate past or potential future experiences (Buzsáki, 2015). SWRs are critical for memory consolidation and retrieval (Buzsáki, 1986; Joo and Frank, 2018), as their disruption impairs spatial learning and memory (Ego-Stengel and Wilson, 2010; Girardeau et al., 2009; Jadhav et al., 2012; Nokia et al., 2012). SWRs are also signatures of a brain-wide activity patterns (Logothetis et al., 2012), and could in principle be detected via EEG or other non-invasive approaches.

We further narrowed our focus to two SWR features that are altered in AD models – SWR abundance and associated slow gamma (SG) power. SWR abundance (events/s) is reduced in apoE4-KI mice (Gillespie et al., 2016) and in models of tau and amyloid β overexpression (Ciupek et al., 2015; Nicole et al., 2016). SWR abundance increases during and after both novel and rewarded experiences (Cheng and Frank, 2008; O’Neill et al., 2008; Singer and Frank, 2009), suggesting a relationship between SWR abundance and the need to store memories. SWR-associated SG power is reduced in both apoE4-KI mice (Gillespie et al., 2016) and in models of amyloid β overexpression (Iaccarino et al., 2016). During SWRs, power in the SG band (30–50 Hz) increases throughout the hippocampus (Carr et al., 2012; Gillespie et al., 2016; Oliva et al., 2018). SWR-associated SG properties have been linked to the quality of memory replay and may help coordinate its structure (Carr et al., 2012; Pfeiffer and Foster, 2015). These observations raise the possibility that SWR abundance and associated SG power could serve as functional biomarkers, but the potential relationship between SWR properties and memory impairments in AD models has yet to be explored. We also know little about the stability of these properties over an animal’s lifetime and about whether SWR properties in early life could predict cognitive abilities in later life.

We set out to determine whether early deficits in SWR features could be used to predict age-dependent cognitive decline in an AD mouse model. We recorded hippocampal network activity from apoE3-KI and apoE4-KI mice at rest, then later examined performance on the Morris water maze (MWM) (Morris, 1984), a spatial goal approach task, and active place avoidance (APA) (Cimadevilla et al., 2000), a spatial goal avoidance task. We found that deficits in SWR abundance and associated SG power in CA3 predicted spatial memory impairment on both tasks in aged apoE4-KI mice. Strikingly, SWR-associated CA3 SG power remained relatively stable over time, and SG power reduction in young apoE4-KI mice predicted spatial memory deficits 10 months later. These findings support the use of SWR properties as functional biomarkers in AD.

## RESULTS

Our study employed a two-stage design. We began with a cohort of animals where we searched for potential predictive relationships between SWR features and behavioral performance in older animals (screen cohort; Figure 1A and Table S1). We then carried out a longitudinal study in a second cohort of animals to determine whether these relationships replicated and whether they were predictive over a substantial fraction of the lifespan (replication cohort; Figure 1A and Table S1).

**Figure 1.**
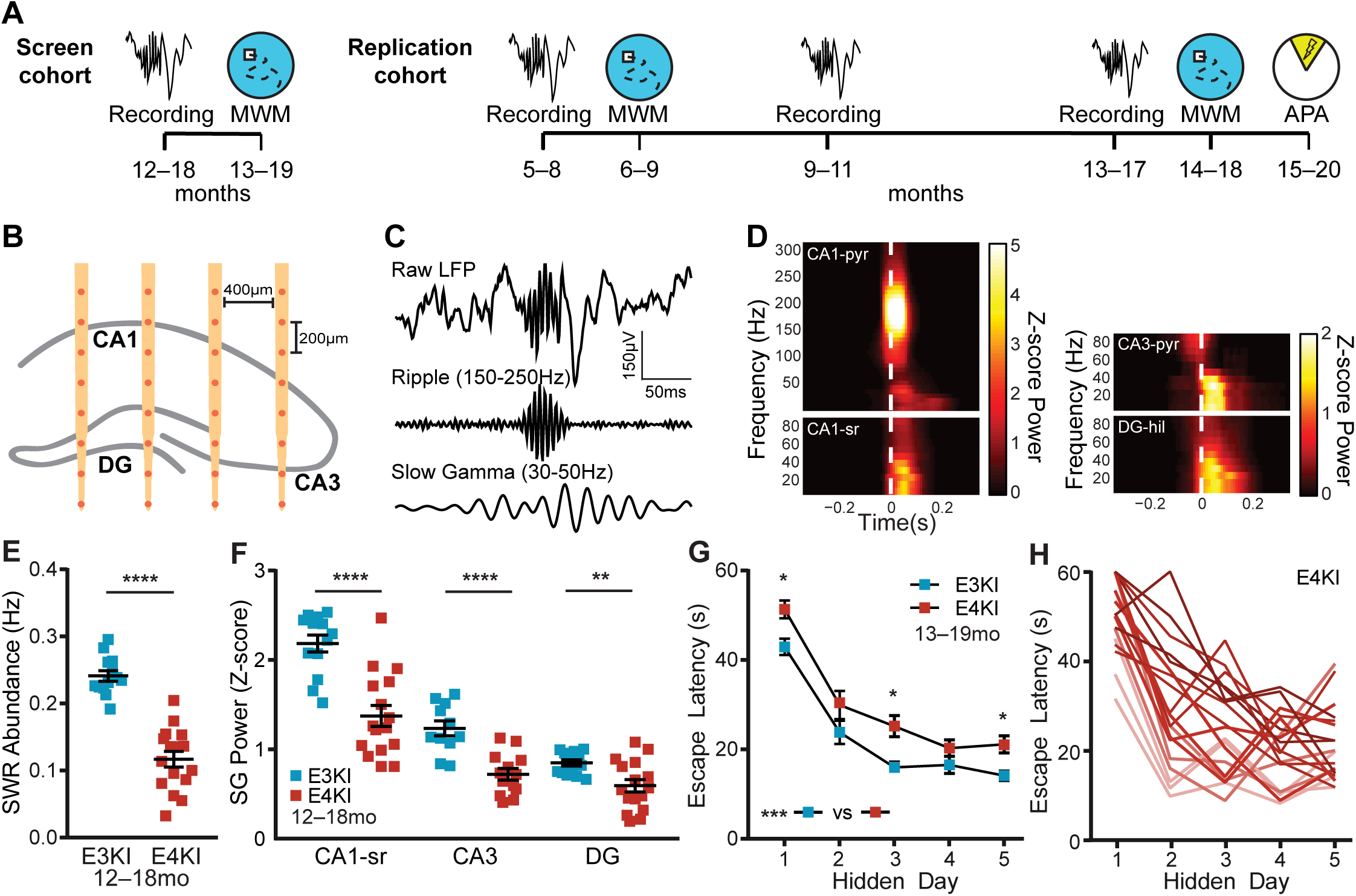
Aged apoE4-KI mice show SWR deficits and spatial approach task impairments and variability. (A) Timeline of experiments. In the screen cohort, aged mice underwent recordings, then MWM 1 month later. In the replication cohort, young mice underwent recordings, then MWM 1 month later, another recording at middle age and old age, then MWM 1 month later and APA 2 months later. (B) Schematic representation of probe placement in dorsal hippocampus, adapted with permission from Gillespie et al., 2016. (C) Representative raw, ripple filtered (150–250Hz), and SG filtered (30–50Hz) traces of a SWR event. (D) Representative SWR-triggered spectrograms from CA1 pyramidal cell layer (CA1-pyr), CA1 stratum radiatum (CA1-sr), CA3 pyramidal cell layer (CA3-pyr), and dentate gyrus hilus (DG-hil) in an apoE3-KI mouse. White dashed lines represent threshold crossing for SWR detection. (E) SWR abundance, n = 13 apoE3-KI and n = 16 apoE4-KI mice, aged 12–18 months (unpaired t test; t(27) = 8.42, p<0.0001). (F) Z-scored SG power during SWRs, n = 13 apoE3-KI and n = 16 apoE4-KI mice (n = 11 and 13 for CA3), aged 12–18 months (unpaired t test, t(27) = 5.17, p<0.0001 for CA1-sr; unpaired t test; t(22) = 4.91, p<0.0001 for CA3; Mann-Whitney U = 45, p = 0.0087 for DG). (G) Average daily escape latency on MWM, n = 20 apoE3-KI and n = 19 apoE4-KI mice, aged 13–19 months. Two-way repeated measures ANOVA of aligned rank transformed data shows significant effect of genotype (F(1,37) = 13.08, p = 0.0009) and post-hoc Mann-Whitney U test with Sidak adjustment shows significant difference on hidden days 1 (U = 95, p = 0.033), 3 (U = 95, p = 0.034), and 5 (U = 92, p = 0.026). (H) ApoE4-KI mice show wide variation in extent of memory impairment. Escape latency curves colored from best (light) to worst (dark) average performance over all days. *p < 0.05; **p < 0.01; ****p < 0.0001. Error bars indicate mean ± SEM. See also Figure S1 and Table S1.

### Aged apoE4-KI mice show SWR deficits and spatial approach task impairments and variability

We first set out to confirm that aged apoE4-KI mice had deficits in SWR features – allowing the use of SWRs as a predictor – and had sufficient individual variation in memory impairment – enabling prediction of this later phenotype. We recorded hippocampal network activity in female apoE3-KI mice and apoE4-KI mice at 12–18 months of age, and then assessed spatial learning and memory on the MWM task one month later (screen cohort; Figure 1A). Recordings were taken from chronically implanted 32-channel silicon arrays targeting right dorsal hippocampus with sites distributed across CA1, CA3, and DG subregions (Figure 1B). Data were collected over five daily 60 min recording sessions in the home cage. SWRs were detected in CA1 stratum pyramidale, and we verified that coincident with these SWRs was an increase in SG power throughout all three subregions of the hippocampus (Figure 1C and 1D; (Gillespie et al., 2016)). One month later, we measured the ability of each mouse to learn the location of a hidden platform in the MWM across 4 daily 1 min trials for 5 days and to remember the previous platform location during 3 probe trials with the platform removed, assessed 24 hours, 72 hours, and 128 hours (probes 1, 2, and 3) after the last hidden platform trial. Replicating previous findings (Andrews-Zwilling et al., 2010; Gillespie et al., 2016), apoE4-KI mice had reduced SWR abundance and associated SG power in CA1, CA3, and DG (Figure 1E and 1F) as well as impaired MWM learning (Figure 1G). Critically, we observed substantial variability in task performance within the apoE4-KI population (Figure 1H) indicating that, just as in human ε4 carriers, genotype is insufficient to explain the extent of cognitive impairment. We capitalized on this variability to explore whether memory impairments could be predicted by deficits in SWR properties.

### SWR deficits predict spatial approach task impairments in aged apoE4-KI mice

Prediction of future outcomes is possible when the value of one variable measured at one time is related to the value of another variable measured at a later time. Prediction can be quantified by measuring the correlation between the two variables. We therefore began our search for a predictive feature by examining the relationship between electrophysiological measurements that showed significant differences between apoE3-KI and apoE4-KI mice and subsequent behavioral performance, focusing on MWM metrics previously shown to be impaired in apoE4-KI mice (Andrews-Zwilling et al., 2010; Gillespie et al., 2016) (Figure 2A). We began with a screen assessing the predictability of 22 behavioral metrics by 4 SWR properties for a total of 88 comparisons, 9 of which were significant with α = 0.05; far more than would be predicted by chance (p < 0.00032, binomial test assuming all initial tests are independent; see Table S2). Furthermore, to ensure that performance metrics were not redundant and captured distinct aspects of behavior, we only included behavior metrics that had an R^2^ < 0.5 with each other throughout this study.

**Figure 2.**
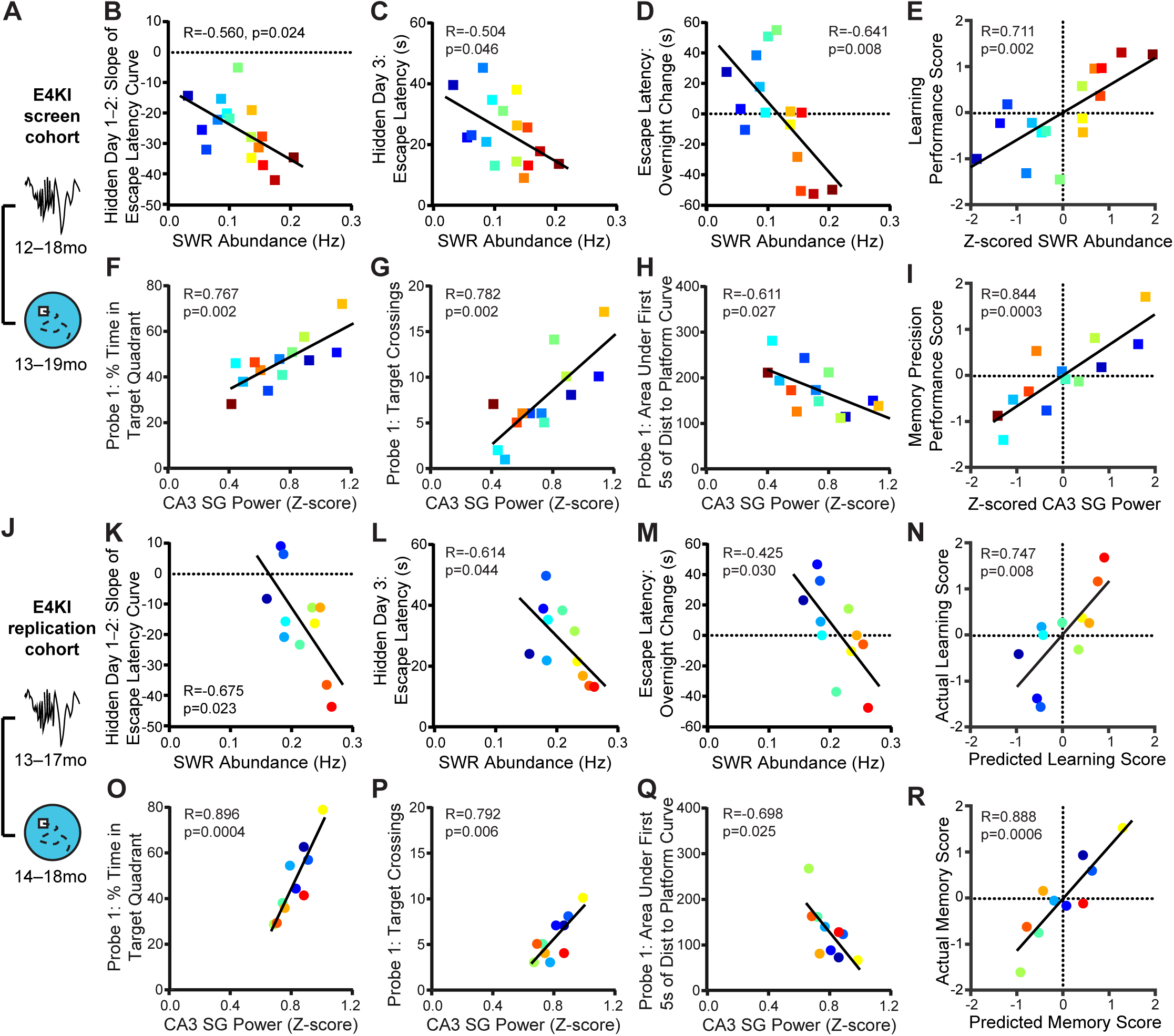
SWR deficits predicts spatial approach task impairments in aged apoE4-KI mice. (A) Timeline of experiments shown in B–I. (B–D) SWR abundance predicts (B) slope of escape latency over hidden days 1–2 (F(1,14) = 6.41), (C) average escape latency on hidden day 3 (F(1,14) = 4.77), and (D) change in escape latency between the last trial of hidden day 1 and the first trial of hidden day 2 (F(1,14) = 9.76), n = 16 mice. (E) Z-scored SWR abundance predicts learning performance score (F(1,14) = 14.3), n = 16 mice. (F–H) CA3 SG power during SWRs predicts (F) percent time spent in quadrant that previously contained the platform (F(1,11) = 15.75), (G) number of times crossing the previous platform location (F(1,11) = 17.3), and (H) area under the curve of the distance to the prior platform location during the first 5 seconds of probe 1 (F(1,11) = 6.55), n = 13 mice. (I) Z-scored CA3 slow gamma power during SWRs predicts memory precision performance score (F(1,11) = 27.3), n = 13 mice. In B–I, apoE4-KI mice aged 12–18 months at electrophysiological recording and 13–19 months at MWM. (J) Timeline of experiments shown in K–R. (K–M) In a replication experiment in a separate cohort of animals, SWR abundance predicts (K) slope of escape latency over hidden days 1–2 (F(1,9) = 7.51), (L) average escape latency on hidden day 3 (F(1,9) = 5.45), and (M) change in escape latency between the last trial of hidden day 1 and the first trial of hidden day 2 (F(1,9) = 6.66), n = 11 mice. (N) Learning performance score as predicted by the linear model in E predicts actual learning performance score (F(1,9) = 11.4), n = 11 mice. (O–Q) In a replication experiment in a separate cohort of animals, CA3 SG power during SWRs predicts (O) percent time spent in quadrant that previously contained the platform (F(1,8) = 32.64), (P) number of times crossing the previous platform location (F(1,8) = 13.48), and (Q) area under the curve of the distance to the prior platform location during the first 5 seconds of probe (F(1,8) = 7.61), n = 10 mice. (R) Memory precision performance score as predicted by the linear model in I predicts actual memory precision performance score (F(1,8) = 30.1), n = 10 mice. In K–R, apoE4-KI mice aged 13–17 months at electrophysiological recording and 14–18 months at MWM. Pearson correlations of apoE4-KI mice. Points colored in order of SWR abundance from blue (lowest) to red (highest). See also Figure S1 and S2.

In aged apoE4-KI mice, SWR abundance (events/s) predicted the slope of the escape latency curve – a measure of the speed of learning – over the first 2 or first 3 hidden days (Figure 2B and S1A). In order to follow individual mice over all measurements, points are colored by SWR abundance from lowest (blue) to highest (red). SWR abundance also predicted escape latency on hidden day 3 (Figure 2C), a measure of approach efficacy. Because SWRs contribute to memory consolidation processes after an experience (Joo and Frank, 2018), we also examined differences between the last trial of a day and the first trial of the next day, between which mice rested for 19 hours (Figure S1B). SWR abundance predicted the extent of behavioral improvement over the first night (Figure 2D), where high SWR abundance predicted improved performance overnight while lower SWR abundance predicted worse performance.

All of these measures relate to rapidity of learning across days, suggesting that lower SWR abundance in aged apoE4-KI mice contributes to slower learning, perhaps through impaired consolidation leading to reduced memory maintenance over days. We therefore combined these four metrics by calculating a z-score for each mouse and metric, reversing the sign of any metrics where lower values indicated better performance, and averaging all z-scores for each mouse. The resulting measure, the learning performance score, is positive to indicate above average performance and negative to indicate below average performance. SWR abundance (also z-scored) accounted for 51% of the variance of the subsequently measured learning performance score (Figure 2E).

Interestingly, SWR-associated SG power in CA3 predicted several metrics of probe memory in aged apoE4-KI mice, but these metrics different from those predicted by SWR abundance. Some mice did not have electrode sites in CA3 and thus were excluded from this analysis (see Table S1). SWR-associated SG power in CA3 predicted the percent time spent exploring the quadrant that previously contained the platform on probes 1 and 2 (Figure 2F and S1C), a measure of retrieval of previously learned spatial information without feedback from the target. To more narrowly define the precision of target location memory, we examined target crossings, which require direct overlap with the previous platform location. CA3 SG power during SWRs predicted the number of target crossings on probe 1 (Figure 2G). Additionally, to examine efficiency of memory retrieval, we measured the cumulative distance traveled toward the previous platform location during the initial approach (first 5 seconds; Figure S1D). CA3 SG power during SWRs predicted this metric for probes 1 and 2 (Figure 2H and S1E). All of these measures relate to precision of memory retrieval, suggesting that lower SWR-associated CA3 SG power in aged apoE4-KI mice reflects an impaired retrieval mechanism. As described above, we combined these 5 metrics, z-scored, into a memory precision performance score. CA3 SG power during SWRs accounted for 77% of the variance in the subsequently measured memory precision score (Figure 2I).

The quantity and strength of these preliminary predictive relationships were compelling enough to motivate a replication and extension of the initial study in a second cohort of animals. Importantly, the analyses of the screen cohort were completed before experiments on the replication cohort, allowing us to establish planned comparisons to determine whether findings from the screen cohort were robust and replicable. For the replication cohort, we implanted young animals (5–8 months) and periodically measured SWR properties in each individual mouse for up to 8 months. We assessed memory performance through MWM tasks at 6–9 months and again at 14–18 months and through an APA task at 15–20 months (Figure 1A and Table S1).

In this replication cohort, studied two years – thus several generations of mice – later, we found that all relationships identified in the screen cohort remained significant (Figure 2J–R and S1F–H). As before, in order to follow individual mice over all measurements, points are colored by SWR abundance from lowest (blue) to highest (red). We also noted that these predictive relationships were consistent across the two cohorts despite differences in the mean values of SWR abundance (µ = 0.12 Hz vs 0.21 Hz; unpaired t test, t(25) = 5.66, p < 0.0001) and the variance of CA3 SG power during SWRs (σ^2^ = 0.05 vs 0.009, F test, F = 5.53, p = 0.015). These differences may have resulted from genetic drift across the multiple generations, colony conditions, or other factors.

Calculating z-scores allowed us to normalize for this difference between the two cohorts, and thus to use the predictive relationships derived from the screen cohort to predict behavioral performance in the replication cohort. These predictions were remarkably accurate, capturing 56% of the variance in the actual learning performance score and 79% of the variance in the actual memory precision performance score (Figure 2N and 2R). Thus, deficits in SWR abundance and associated CA3 SG power, z-scored across the population, predict early learning and memory precision impairment, respectively, even when applied to a separate cohort of animals.

These predictive relationships were also robust: we determined that none of the correlations in this study were driven by a single data point (see STAR Methods). Further, we established in the replication cohort that there were no genotype differences in anxiety or exploratory drive as measured by open field and elevated plus maze tests, in pain response as measured by a hot plate test, or in visual acuity as measured by MWM trials with the platform marked (Figure S2). Moreover, none of these non-spatial behaviors significantly correlated with spatial task performance or with SWR properties at α = 0.05. Therefore, spatial performance differences were most likely driven by spatial memory ability, and SWR-related network alterations were specifically related to this spatial memory impairment. Finally, while we identified strong and replicable predictive relationships between SWR properties and memory for aged apoE4-KI mice, no such relationships were observed for aged apoE3-KI mice in either cohort. This may be the result of a ceiling effect given the overall much higher levels of both SWR abundance and associated CA3 SG power in apoE3-KI mice (Figure 1E and 1F; for example see Figure S1I). Additionally, SWR-associated SG power in CA1 and in DG did not consistently and significantly correlate with behavior, thus we focused only on SWR abundance and associated CA3 SG power for the rest of the study.

### SWR deficits predict spatial avoidance task impairments in aged apoE4-KI mice

A robust predictor of memory decline should generalize across tasks, so we next assessed the predictive capacity of SWR properties for subsequent performance on the APA task (Figure 3A), a spatial avoidance task (Cimadevilla et al., 2000). In this task, mice explore a rotating arena and must use distal cues to avoid a shock zone that is fixed relative to the room across daily 10 min trials for 4 days. As this task has not been previously used in AD models, we first asked whether there was an overall effect of apoE genotype. ApoE4-KI mice task acquisition was significantly impaired on day 1; mice had greater number of entrances into the shock zone (Figure 3B), travelled less distance (Figure S3A), spent less time in the quadrant opposite the shock zone (Figure S3B), and moved further from the shock zone in bouts of movement (Figure S3C). Performance on the probe trial, in which the shock was inactivated, was also impaired (Figure S3D). Thus, APA is able to identify significant behavioral differences between aged apoE4-KI and apoE3-KI mice.

**Figure 3.**
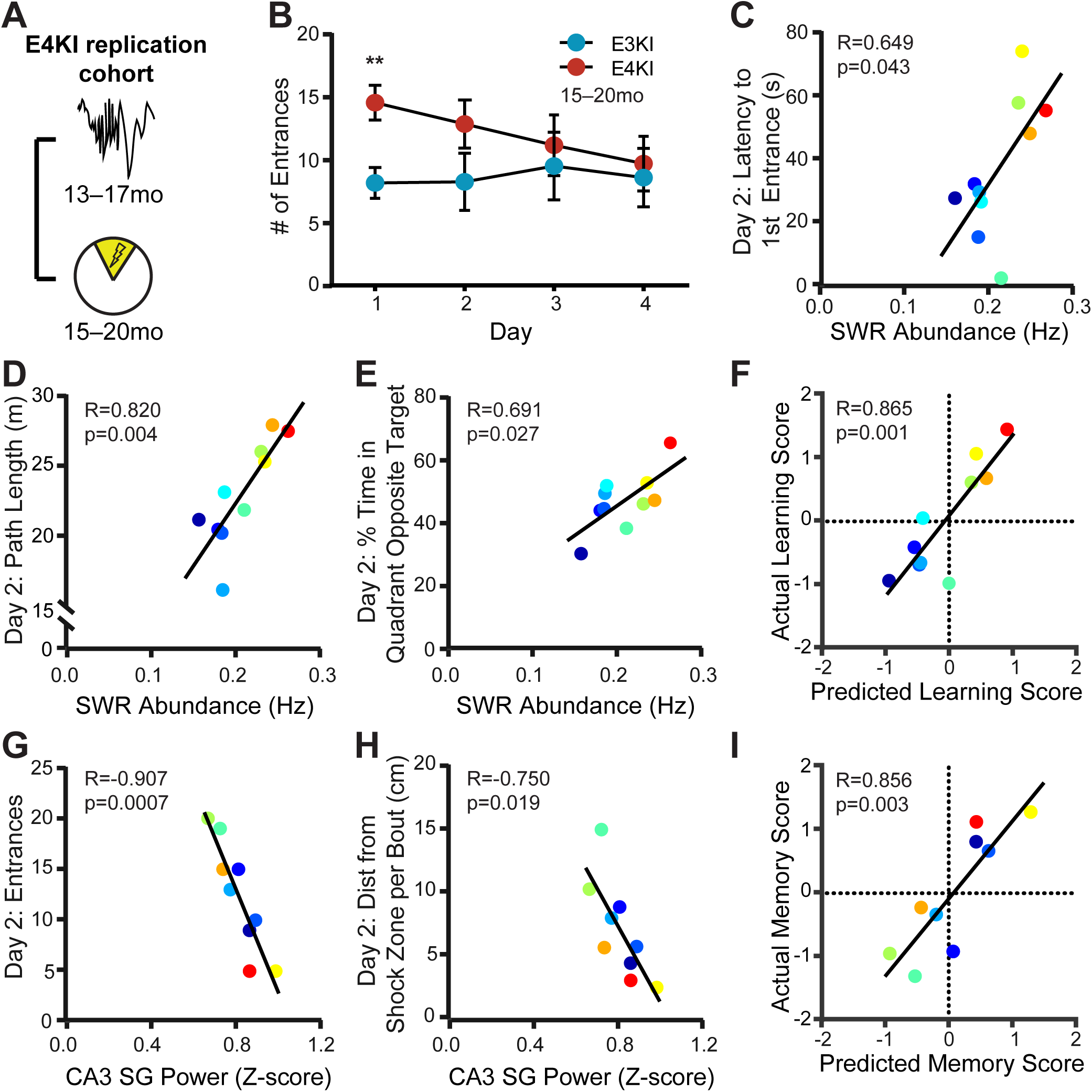
SWR deficits predict spatial avoidance task impairments in aged apoE4-KI mice. (A) Timeline of experiments shown in B–I. (B) Number of entrances into the shock zone is significantly different on day 1 (unpaired t test with Sidak’s multiple comparison adjustment, t(23) = 3.43, p = 0.009). n = 12 apoE3-KI mice and n = 13 apoE4-KI mice, aged 15–20 months. **p < 0.01. Error bars indicate mean ± SEM. (C–E) SWR abundance predicts (C) latency to first shock zone entrance (F(1,8) = 5.81), (D) total path length (F(1,8) = 16.42), and (E) percent of total time spent in quadrant opposite the shock zone (F(1,8) = 7.33) on day 2, n = 10 apoE4-KI mice. (F) Learning performance score as predicted by the linear model in 2E predicts actual learning performance (F(1,8) = 23.7), n = 10 apoE4-KI mice. (G,H) CA3 SG power during SWRs predicts (G) entrances to the shock zone on day 2 (F(1,7) = 32.53) and (H) distance travelled per movement bout relative to shock zone boundaries on day 2 (F(1,7) = 9.04), n = 9 apoE4-KI mice. (I) Memory precision performance score as predicted by the linear model in 2I predicts actual memory precision performance score (F(1,7) = 19.2), n = 9 apoE4-KI mice. Mice aged 15–20 months at APA and 13–17 months at electrophysiological recording. Points colored in order of SWR abundance from blue (lowest) to red (highest). All correlations are Pearson correlations of apoE4-KI mice. See also Figure S2 and S3 and Table S3.

We then determined which behavioral metrics correlated with SWR properties, noting that, as in MWM, there was substantial variability in apoE4-KI mice (Figure S3E). In total, we compared 5 behavioral metrics over 4 days against 2 SWR properties for a total of 40 comparisons, 8 of which were significant with α = 0.05; far more than would be predicted by chance (p < 0.00071, binomial test assuming all initial tests are independent; see Table S3). Most of the significant correlations were found on day 2. We computed the variance in the number of shock zone entrances across animals for days 1 and 2 and found that day 2 variances were higher (mean entrances day 1 σ^2^ = 15.76 vs day 2 σ^2^ = 33.82 mean entrances), suggesting day 2 performance would be more effective for assessing predictive validity. In addition, day 2 performance predicted day 3 and 4 performance, and apoE4-KI mice separated out into two distinct performance populations on day 2 such that the top 50% of performers on day 2 continued to learn on days 3 and 4 and that the bottom 50% did not (Figure S3F and S3G), suggesting that predicting day 2 performance could capture much of the learning curve.

SWR abundance measured 2 months before task training predicted the latency to first entrance on day 2, a measure of memory of the shock location assessed 24 hours after the previous training trial and before receiving any feedback in that trial (Figure 3C). This result closely parallels the measure for overnight consolidation in the MWM task, which also correlated with SWR abundance (Figure 2D and 2M). SWR abundance also predicted path length and percent time in the quadrant opposite the target on day 2 (Figure 3D and 3E). Together, these findings suggest that SWR abundance deficits in apoE4-KI mice are related to avoidance efficacy, just as it correlated with approach efficacy on hidden day 3 of the MWM (Figure 2C and 2L). We therefore calculated a learning performance score for the APA task in the same manner as for the MWM task (mean for each animal across the z-scored metrics shown in Figure 3D and 3E) and used the predictive relationship derived from the screen cohort to assess our ability to predict relative behavioral impairments from SWR abundance. Strikingly, the relationship between the predicted and actual learning performance scores was very strong, with the predicted score accounting for 75% of the variance in the actual score (Figure 3F).

A similar pattern of predictability was seen for CA3 SG power during SWRs. This SWR property predicted the number of shock zone entrances – a metric which requires precise memory of shock zone boundaries – on day 2 (Figure 3G). CA3 SG power during SWRs also predicted the number of shock zone entrances for days 3 and 4, and thus captured performance beyond early learning, but these were excluded from further consideration as these metrics were highly correlated across days (Figure S3F). CA3 SG power during SWRs further predicted the distance mice travelled away from the shock zone during bouts of movement on days 2 and 3 (Figure 3H and S3H). Mice with lower CA3 SG power during SWRs moved further from the shock zone, suggesting less precise memory of shock zone boundaries. This is consistent with the MWM results, in which SWR-associated CA3 SG power deficits in aged apoE4-KI mice are related to impairments in memory retrieval precision, particularly later in the task (Figure 2F–H and 2O–Q).

As above, we calculated a memory precision performance score for the APA task in the same manner as for the MWM task (mean for each animal across the sign reversed z-scored metrics shown in Figure 3G, 3H, and S3H) and predicted memory precision performance scores using the model developed in the screen cohort. Here again, the relationship between the predicted and actual memory precision performance scores was very strong, with the predicted score accounting for 73% of the variance in the actual score (Figure 3I). Thus, the models developed in the screen cohort accurately predict behavioral performance, despite being derived from a different behavior and a different group of animals. We further noted that the performance scores of the MWM task predicted the performance scores of the APA task (Figure S3I and S3J). Therefore, performance on the first 3 days of MWM training predicted avoidance efficacy on day 2 of APA, while performance on probe trials during MWM predicted memory precision on day 2 of APA.

### SWR deficits at younger ages predict spatial approach and spatial avoidance task impairments at older ages

Finally, a meaningful predictor should be consistent within a single subject over aging and have predictive power before the onset of memory impairment. We conducted a longitudinal study of the replication cohort, measuring behavior and electrophysiology from the same mice at 5–8 months (young), 9–11 months (middle-aged), and 13–17 months (old) (Figure 1A, 4A, and Table S1). Young apoE4-KI mice already had reduced SWR abundance and associated SG power in CA3 when compared to young apoE3-KI mice (Figure S4A and S4B). Interestingly, SWR abundance increased in apoE3-KI mice over aging, although there was also a decrease in SWR baseline power (Figure S4C) that, along with a preserved standard deviation of SWR power (Figure S4D), could have contributed to this increase. Interestingly, increases in CA3 SG power during SWRs were seen in both groups over aging (Figure S4B) without concomitant changes in baseline or standard deviation (Figure S4E and S4F), suggesting SWR-specific physiological changes and indicating stability in the quality of the recordings.

Overall, these findings suggest that these SWR properties, rather than degrading with age, may already show deficits at young ages that will manifest as behavioral deficits later, facilitating their potential use as an early predictor for later memory decline. The early reduction in SWR abundance and associated SG power in CA3 did not translate into an early behavior deficit however: young apoE4-KI mice showed no detectable learning deficits (Figure S4G) (Leung et al., 2012), and SWR abundance and associated CA3 SG power measured at 5–8 months did not predict behavior tested one month later (Table S4). The lack of early memory deficits in the presence of altered network activity related to spatial navigation parallels observations young adult human ε4 carriers (Kunz et al., 2015) and suggests the possibility of compensation in the younger brain.

We then asked whether SWR properties were consistent within individual mice across time. We found that while SWR abundance was not significantly correlated across ages in either apoE3-KI or apoE4-KI mice (Figure 4B and 4C), CA3 SG power during SWRs was significantly correlated across ages in both genotypes (Figure 4D and 4E). We noted that baseline power in the SWR frequency band, but not in the SG frequency band, declined over aging, which could explain this lack of correlation for SWR abundance. Despite the significant prediction over 4 months in CA3 SG power, this did not hold true over longer time scales. CA3 SG power at 5–8 months did not significantly predict CA3 SG power at 13–17 months (e.g. for apoE3-KI mice, R = 0.2, F(1,12) = 0.5, p = 0.49), thus CA3 SG power during SWRs was unlikely to predict the same behavioral metrics when measured at 5–8 months as when measured at 13–17 months. Overall, these findings indicate that CA3 SG power during SWRs is a relatively stable measure over the timescale of four months, making it a more promising candidate for a predictive measure over aging.

**Figure 4.**
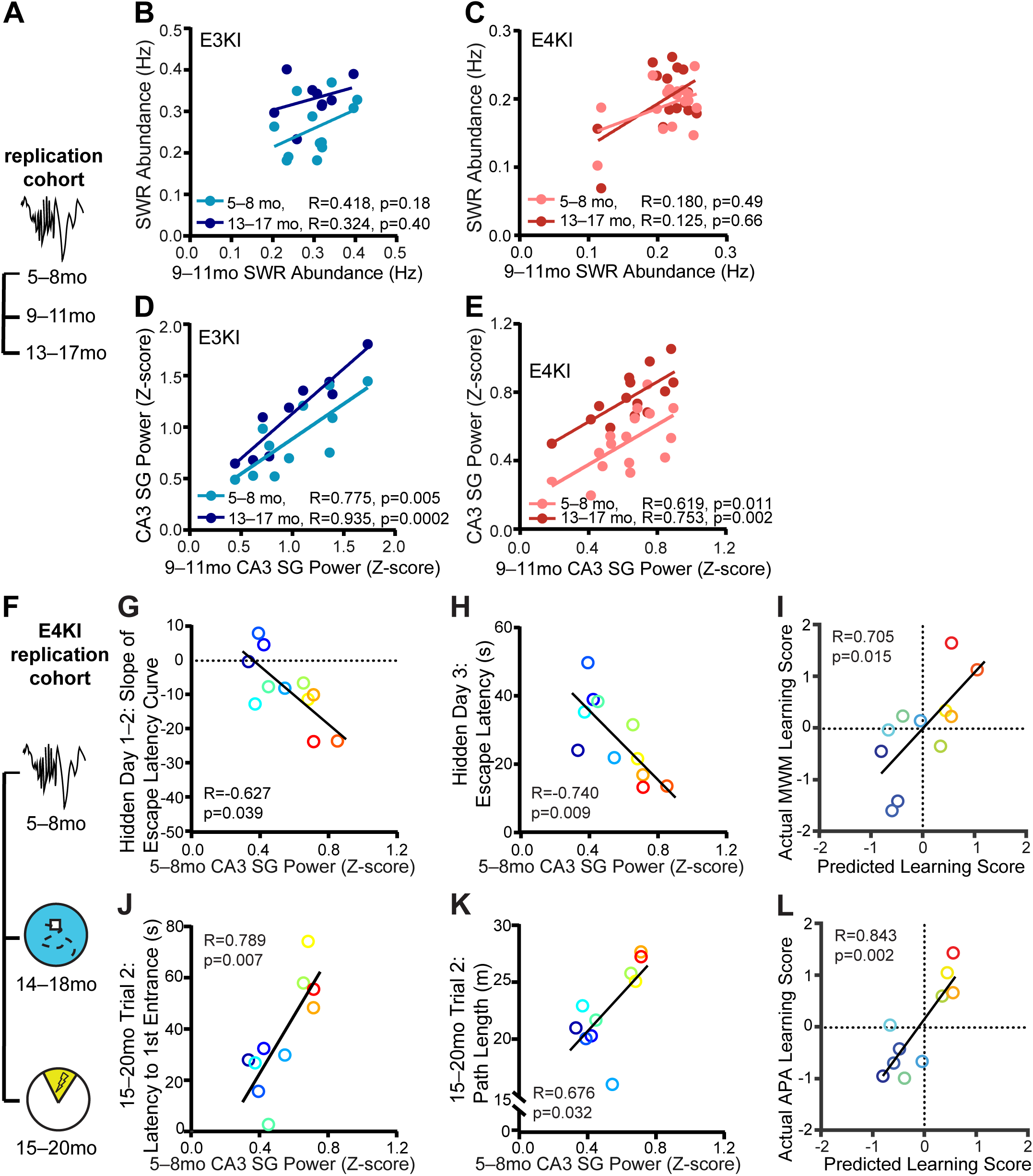
SWR deficits at younger ages predict spatial approach and spatial avoidance task impairments at older ages. (A) Timeline of experiments shown in B–E. (B,C) SWR abundance at 9–11 months does not correlate with SWR abundance at 5–8 months and 13–17 months in (B) apoE3-KI mice (F(1,10) = 2.12, n = 12 for 5–8 months, F(1,7) = 0.82, n = 9 for 13–17 months) and (C) apoE4-KI mice (n = 17 for 5–8 months, n = 15 for 13–17 months, Spearman correlation). (D,E) CA3 SG power during SWRs at 9–11 months correlates with CA3 SG power during SWRs at 5–8 months and 13–17 months in (D) apoE3-KI mice (F(1,9) = 13.54, n = 11 for 5–8 months, F(1,7) = 48.52, n = 9 for 13–17 months) and (E) apoE4-KI mice (F(1,14) = 8.71, n = 16 for 5–8 months, F(1,12) = 15.71, n = 14 for 13–17 months). (F) Timeline of experiments shown in G–L. (G,H) CA3 SG power during SWRs measured at 5–8 months predicts (G) slope of escape latency over hidden days 1–2 (F(1,9) = 5.82) and (H) average escape latency on hidden day 3 (F(1,9) = 11.08) on MWM task at 14–18 months, n = 11 apoE4-KI mice. (I) MWM learning performance score as predicted by the linear model in 2E predicts actual learning performance score (F(1,8) = 8.9), n = 11 apoE4-KI mice aged 5–8 months at electrophysiological recording and 14–18 months at MWM. (J,K) CA3 SG power during SWRs also predicts (J) latency to first shock zone entrance (F(1,8) = 13.21) and (K) total path length (F(1,8) = 6.74) on APA task at 15–20 months, n = 10 apoE4-KI mice. (L) APA learning performance score as predicted by the linear model in 2E predicts actual learning performance score (F(1,8) = 19.7), n = 10 apoE4-KI mice aged 5–8 months at electrophysiological recording and 15–20 months at APA. Points colored in order of 13–17 month SWR abundance from blue (lowest) to red (highest). Pearson correlations unless otherwise specified. See also Figure S4 and Table S4.

Indeed, when we examined the relationship between CA3 SG power during SWRs and behavior 10–11 months later (Figure 4F), we found strong predictive relationships (Table S4). CA3 SG power during SWRs measured at 5–8 months predicted escape latency on the third hidden day and the slope of the escape latency curve for the first 2 or first 3 days on the MWM task at 14– 18 months (Figure 4G, 4H, S4H, and Table S4). It also predicted the number of entries into the shock zone and path length on day 2 during the APA task at 15–20 months (Figure 4J, 4K and Table S4). As before, points are colored by SWR abundance measured at 13–17 months from lowest (blue) to highest (red). Therefore, deficits in CA3 SG power during SWRs in young apoE4- KI mice – before spatial learning impairment is detectable – predicted future learning impairment on both a spatial approach and a spatial avoidance task at older ages. We confirmed that these effects were not expected given the multiple comparisons made: in total, we compared 15 behavioral metrics against 2 SWR properties for a total of 30 comparisons, 5 of which were significant with α = 0.05; far more than would be predicted by chance (p < 0.016, binomial test assuming all initial tests are independent; see Table S4).

Of all behavioral measures related to learning as established in Figures 2 and 3, 3 of 4 MWM metrics and 2 of 3 APA metrics were significantly predicted by CA3 SG power during SWRs measured at 5–8 months. To assess our ability to make predictions of overall behavioral performance based on the relationships in the screen cohort, we once again combined all 4 MWM metrics into a learning performance score (Figure 4I) and all 3 APA metrics into an APA learning performance score (Figure 4L). Here again, the relationship derived from the screen cohort effectively predicted behavioral deficits, allowing us to account for 50% and 71% of the variance in the actual learning performance score with CA3 SG power during SWRs measured 10–11 months previously.

We were surprised to note that the behavioral measures that could be predicted by SWR abundance measured at older ages were predicted by SWR-associated CA3 SG power measured at young ages. This suggests a relationship between CA3 SG power at 5–8 months and SWR abundance measured at 13–17 months, and indeed CA3 SG power during SWRs in young apoE4- KI mice strongly predicted SWR abundance 8 months later (Figure S4I). This was not true for young apoE3-KI mice, again perhaps due to a ceiling effect (Figure S4I). Therefore, in apoE4-KI mice, the amount of SG power generated in CA3 during SWRs in a young mouse predicted the extent to which that mouse generated SWRs at rest 8 months later.

## DISCUSSION

We have demonstrated that SWR abundance and associated CA3 SG power can be used as functional biomarkers to predict cognitive decline in an apoE4 mouse model of AD. We capitalized on the behavioral population variability and the SWR-related network alterations found in apoE4- KI mice, which allowed us to observe correlations between the two. We then observed that these SWR properties, measured 1–2 months prior to behavior in aged apoE4-KI mice, predict multiple behavioral metrics, capturing different but related aspects of spatial learning and memory. These findings were not restricted to a single cohort, as demonstrated through replication, or to a single task, as demonstrated through correlations observed with two distinct spatial tasks. Finally and most critically, CA3 SG power during SWRs in young apoE4-KI mice, measured prior to the onset of detectable cognitive deficits, predicted spatial learning and memory impairments across both tasks 10 months later. This metric was also correlated within individual animals over several months, making it a potential functional biomarker for cognitive decline in AD.

We observed differences in both SWR abundance and associated CA3 SG power between the screen and replication cohorts. While experimental conditions were identical to the greatest practical extent, it is possible that genetic drift occurred across multiple generations or that colony conditions changed, which could affect this result. Despite this, when we z-score normalized the data, we found that predictive relationships derived from the initial screen cohort were very strongly predictive in the replication cohort for both the MWM and the APA tasks. This consistence suggests that a simple normalization to the values seen in a given population allows for accurate predictions of behavioral impairments from physiological measurements across many months in our mouse models.

Through chronic measurement of hippocampal network activity in individual mice, we were able to assess whether SWR features are stable over several months. SWR abundance is affected by environmental variables such as novelty and reward (Cheng and Frank, 2008; Singer and Frank, 2009), but such a relationship has yet to be defined for SWR-associated slow gamma. Our results suggest that over timescales of 4 months, CA3 SG power during SWRs is significantly correlated with itself, while SWR abundance is not, perhaps due to changes in baseline activity in the SWR frequency band over aging. Previous cross-sectional work in wild type rats reported that SWR abundance measured during or immediately after a task was reduced in aged rats (Cowen et al., 2018; Wiegand et al., 2016). Our longitudinal measurements in human apoE3-KI mice during rest, independent from task performance, did not show this reduction. Rather, we observed that both SWR abundance and associated CA3 SG power slightly increased with aging, although average measures for apoE4-KI mice remained lower than for apoE3-KI mice across all ages. Since behavioral differences do not emerge until later ages, this suggests an additional network change – perhaps a loss of compensation – that occurs over aging in apoE4-KI mice, which interacts with existing physiological deficits to cause behavioral impairment.

Notably, we did not observe any consistent and significant correlations between CA1 or DG SG power during SWRs and spatial cognitive performance. Substantial previous work has shown that physiological characteristics of CA3 can distinguish between aged rats with impaired or unimpaired memory. Aged impaired rats have elevated CA3 activation, which leads to inability to remap CA3 place cells in novel contexts (Haberman et al., 2017; Wilson et al., 2005). They furthermore have reduced expression of genes related to synaptic plasticity in CA3 (Adams et al., 2001; Haberman et al., 2011, 2013; Smith et al., 2000). Our findings establish another way in which measurements from CA3 can distinguish impaired from unimpaired spatial memory. Moreover, SG activity throughout the hippocampus is hypothesized to originate from CA3. During SWRs, CA1 SG is most coherent with CA3 SG activity, and SG power is highest in the stratum radiatum, the input layer from CA3 (Carr et al., 2012; Gillespie et al., 2016; Ramirez-Villegas et al., 2018). Thus, it is reasonable that SG power measured at its hypothesized generator is the strongest predictor of spatial learning and memory.

The extent of coherence between CA1 and CA3 of oscillations in the SG frequency band correlates with replay fidelity, the temporal order of place cell firing in a spatial sequence (Carr et al., 2012). Notably, one previous study found that measures of replay fidelity following a linear track run correlated with MWM learning in aged wild type rats (Gerrard et al., 2008). Together with our finding of correlations between CA3 SG power during SWRs and MWM performance, this suggests that measures related to replay fidelity measured outside of task performance can predict MWM learning deficits in the context of both normal aging and AD aging.

We additionally found that CA3 slow gamma power during SWRs at young ages predicts SWR abundance 8 months later. While the cause of this correlation is unclear, it suggests a relationship in apoE4-KI mice between organization of replay events in early adulthood and the number of events later in life. There may be a positive feedback loop in which replays with greater fidelity to the original encoded sequences lead to greater probability of future replays over aging. Alternatively, CA3 SG power during SWRs may be a more sensitive reflection of the health of the underlying circuitry, which may affect SWR generation in later life.

The most common proposed biomarkers for AD are amyloid or tau, measured in CSF or by PET imaging (Riedel, 2014). However, at least 40% of cognitively normal elderly patients show amyloid or tau pathology, indicating that these molecular biomarkers are not sufficient to distinguish healthy aging from AD-induced cognitive decline (Boyle et al., 2013; Caselli and Reiman, 2013). Hippocampal network activity shows a clear link between pathology and behavioral outcomes and represents a potential new class of functional biomarkers. Moreover, human longitudinal studies of potential biomarkers have only been able to follow sporadic AD subjects for 3–8 years prior to diagnosis. Since cognitive decline accelerates 15-fold during the 5– 6 years prior to AD diagnosis, these biomarkers may only be sufficient to predict ongoing cognitive decline (Wilson et al., 2011). In contrast, this study in mice was able to predict cognitive decline long before memory impairment was detectable.

A physiological biomarker also has the distinct advantage of being compatible with repeated measures. In contrast, behavioral readouts often cannot be repeated without affecting performance, and post-mortem pathology only captures the final endpoint. Using a network signature, disease progression could be measured over aging, and cellular pathological differences that differentiate future impaired from unimpaired animals could be assessed at an early age. During preclinical drug studies, a single animal could be measured over a time course of treatment or with variable doses, both reducing the number of animals required and increasing study power through use of paired statistics. Most critically, preventative therapies could be tested in animals by measuring the effect on the hippocampal network before spatial learning and memory impairments arise.

Currently, SWRs are only detectable in humans by depth electrodes (Axmacher et al., 2008), thus new technology will need to be developed to translate this functional biomarker into humans. However, SWRs are signatures of a brain-wide activity patterns (Logothetis et al., 2012), and could in principle be detected via EEG or other non-invasive approaches. In particular, recently MEG has been demonstrated to be capable of detecting oscillations in the same frequency range in deep brain structures (Yin et al., 2019), and high density scalp EEG has been shown to detect signals highly correlated to those measured by intracranial electrodes in deep brain structures (Seeber et al., 2019). Moreover, recent work in humans shows that SWRs detected in the medial temporal lobe are highly correlated with SWRs detected in the temporal association cortices, suggesting that detecting SWRs from cortex could serve as a proxy (Vaz et al., 2019). Overall, SWR features are compelling functional biomarker candidates that can predict future cognitive decline in an apoE4 model of AD and could potentially be used to predict AD risk and assess treatment efficacy before the onset of symptoms.

## ACKNOWLEDGEMENTS

This work was supported by the National Science Foundation Graduate Research Fellowship No. 1144247, National Institute on Aging Predoctoral Fellowship No. F31AG057150, and Genentech Foundation Fellowship to E.A.J, a Simons Collaboration for the Global Brain Fellowship to A.K.G., funding from the Howard Hughes Medical Institutes to L.M.F., and National Institute on Aging grants RF1AG047655, RF1AG055421, and R01AG055682 to Y.H. We thank Desiree Macchia and T. Michael Gill for assistance with the place avoidance task, Hanci Lei and Alyssa Yang for assistance with histology, and Brian Grone, Marielena Sosa, and Kelly Zalocusky for feedback on the manuscript.

## AUTHOR CONTRIBUTIONS

E.A.J., A.K.G., and Y.H. designed and coordinated the study. E.A.J. carried out most studies and data analysis and wrote the manuscript. A.K.G. contributed to the studies and analysis of the screen cohort. S.Y.Y. managed mouse lines, performed perfusions, and conducted the MWM tests. A.K.G., L.M.F., and Y.H. provided advice on data analysis and interpretations and edited the manuscript. L.M.F. and Y.H. supervised the project.

## COMPETING FINANCIAL INTERESTS

Y.H. is a co-founder and scientific advisory board member of E-Scape Bio, Inc. and GABAeron, Inc. Other authors declare no competing financial interests.

## STAR METHODS

### Contact for Reagent and Resource Sharing

Further information and requests for resources and reagents should be directed to and will be fulfilled by the Lead Contact, Yadong Huang.

### Experimental Model and Subject Details

#### Mice

Mice with human apoE3 or apoE4 knocked-in at the mouse *APOE* locus on a C57BL/6 background (Sullivan et al., 2004) were originally obtained from Taconic. All animals were bred in-house using trio breeding producing 10 pups per litter on average, which were weaned at 28 days. Female mice aged 5–20 months were used. Animals were housed in a pathogen-free barrier facility on a 12h light cycle (lights on at 7am and off at 7pm) at 19–23°C and 30–70% humidity. Animals were identified by ear punch under brief isofluorane anesthesia and genotyped by PCR of a tail clipping at both weaning and perfusion. All animals otherwise received no procedures except those reported in this study. Throughout the study, mice were singly housed. All animal experiments were conducted in accordance with the guidelines and regulations of the National Institutes of Health, the University of California, and the Gladstone Institutes under IACUC protocol AN117112.

### Method Details

The study consisted of two cohorts of female apoE3-KI and apoE4-KI mice: one screen cohort which had electrophysiological recordings at 12–18 months and MWM at 13–19 months and one replication cohort which had electrophysiological recordings at 5–8 months, 9–11 months, and 13– 17 months, MWM at 5-8 months, open field, elevated plus maze, and MWM at 14–18 months, and APA and hot plate at 15–20 months (see Figure 1A). Electrode lifetime did not diminish our ability to detect SWRs and measure associated SG power, as neither metric declined over aging (Figure S4A and S4B). Some mice in the replication cohort did not survive the duration of the longitudinal study, so their electrophysiological data was included in younger group analyses, but their behavioral data was not available for group analyses or correlations (Table S1). For this reason, additional mice which had not had any LFP recordings were included in the aged behavioral studies of the replication cohort to achieve sufficient power for group analyses (Table S1). All procedures were conducted during the light cycle. Mice were recorded at a randomly allocated time each day to counteract differences caused by circadian effects. The experimenters were blinded to genotype during surgery, recordings, behavior, and histology.

#### Surgery

Mice were anesthetized by intraperitoneal injection of ketamine (60 mg/kg) and xylazine (30 mg/kg); anesthesia was maintained with 0.6–1.5% isofluorane given through a vaporizer and nose cone. The head was secured with earbars and a tooth bar in a stereotaxic alignment system (Kopf Instruments). Fur was removed from the scalp, which was then sterilized with alternating swabs of chlorhexidine and 70% ethanol. The scalp was opened, sterilized with 3% hydrogen peroxide, and thoroughly cleaned to reduce risk of tissue regrowth. 0.5 mm craniotomies were made over the right frontal cortex and left parietal cortex. Skull screws (FST) were inserted to anchor and support the implant, and were secured with dental adhesive (C&B Metabond, Parkell). An additional 0.5 mm craniotomy was made over the right cerebellum for insertion of the indifferent ground and reference wires. A forth craniotomy was centered at −1.95 mm AP and 1.5 mm ML from bregma and extended bidirectionally along the ML axis to 2 mm width to receive the recording probe. The probes had four 5 mm shanks spaced 400 µm apart with 8 electrode sites per shank and 200 µm spacing between sites (Neuronexus; configuration A4×8-400-200-704-CM32). The probe was quickly lowered until the tip reached 2.2 mm below the surface of the brain, and the reference and ground wire was inserted into the subdural space above the cerebellum. The probe was cemented in place with dental acrylic and the scalp was closed with nylon sutures. Mice were treated with 0.0375 mg/kg buprenorphine intraperitoneally and 5 mg/kg ketofen subcutaneously 30–45 min after surgery, monitored until ambulatory, then monitored daily for 3 days. A minimum of 1 week was allowed for recovery before recording.

#### Electrophysiology

Data from all mice was collected, amplified, multiplexed, processed, and digitized with 32-channel Upright Headstage, commutator, and Main Control Unit (SpikeGadgets). Simultaneous data acquisition at 30 kHz and video tracking at 30 frames/s was performed using Trodes software (SpikeGadgets). Each data collection time point consisted of 5 days of 60 min home cage sessions. Each mouse was recorded at a randomly assigned time each day across the light circle to control for the effects of circadian rhythm. During recordings, home cages were changed to Alpha-dri bedding (Shepherd Specialty Papers) to enable video tracking.

#### Behavior

During the MWM task, mice were housed in the testing room with the arena obscured by a partition and given 2 days to acclimate to the room in a fresh cage before training began. In all trials, mice were placed in a 122 cm diameter pool filled with opaque water and had to locate a 14 × 14 cm platform submerged 1.5 cm below the water’s surface. Mice could only use spatial cues on the walls around the pool to guide their search. On once daily pretraining trial for the first 2 days, mice swam down a rectangular channel until they locate the platform, or were guided there by the experimenter after 90 s. Then, the rectangular guides were removed and the platform was placed in a new location. On 4 daily trials for the next 5 hidden days, mice were dropped at random locations each trial and given 60 s to locate the platform. Daily trials were divided into two pairs 10 min apart, with 4 hours between the pairs. Then, on probe trials conducted 24 hours, 72 hours, and 128 hours after the last hidden day, the platform was removed, and mice explored the arena for 60 s. Finally, on twice daily visible trials over 3 days, a flag was placed on the platform, and mice swam directly to the platform to measure visual acuity. Video tracking at 30 frames/s was performed during all trials with Ethovision (Noldus).

During the APA task, mice were housed adjacent to the testing room and given 3 days to acclimate to the room in a fresh cage before training began. In all trials, mice were placed in a 40 cm diameter arena that rotates at 1 rpm (BioSignal Group). On the first day, mice habituated to the environment by exploring it for 10 minutes. On the following 4 days, when mice entered a 60° region which is fixed relative to the room, as measured by video tracking, they received a 0.2 mA foot shock for 500 ms at 60 Hz every 1.5 s until they left the shock zone. Mice could only use spatial cues on the walls around the arena to actively avoid this shock zone given the constant rotation of the arena. On the fifth day, the shock was inactivated for the first 5 min (probe), then turned on for the second 5 min (reinstatement) of a 10 min continuous trial. 1 apoE4-KI mouse was excluded from APA and hot plate testing due to a motor impairment it developed immediately prior to testing (see Table S1). Shock times and video tracking at 30 frames/s relative to the rotating arena were recorded during all trials with Tracker software (BioSignal Group).

We selected five metrics to assess different aspects of spatial learning and memory on the APA task, four of which were proposed by the original task creators (Cimadevilla et al., 2000). Over a trial, mice attempt to enter the shock zone as few times as possible (entrances). Mice are successful if they move around in the arena to avoid the shock zone (path length) rather than staying still and then fleeing when shocked, optimally spending most of the trial as far from the shock zone as possible (percent time in quadrant opposite target). Mice with more accurate representations of the shock zone boundary may not move far from the shock zone during each bout of movement (distance from shock zone per bout). At trial start, mice avoid the shock zone before being given any feedback as to its location (latency to first entrance).

Three tasks were used to assess general anxiety, exploratory drive, and nocioception in order to determine if these factors affected spatial memory task acquisition. For these tests, mice were habituated to the testing room for 1 hour prior to testing. First, during the open field test, mice explore a 41 cm x 41 cm enclosed arena (San Diego Instruments). Location and movements are captured by beam breaks and analyzed in Photobeam Activity System software (San Diego Instruments). Reduced time spent or distance travelled in center of the field indicates anxiety, and reduced total distance travelled indicates reduced exploratory drive. Second, during the elevated plus maze, mice explore a plus-shaped maze 80 cm above the ground, 5 cm in width and 75 cm in length, consisting of 2 open arms without walls and 2 closed arms with 17 cm walls (Kinder Scientific). Location is captured by beam breaks and analyzed in MotorMonitor software (Kinder Scientific). Reduced time spent or distance travelled in open arm indicates anxiety, and reduced total distance travelled indicates reduced exploratory drive. Third, during the hot plate test, mice are placed on a 52°C plate in an open cylinder (Campden Instruments). Latency to hindpaw withdrawal is recorded via live observation by the experimenter, with high latency suggesting impaired nocioception.

#### Histology

Mice were deeply anesthetized with avertin, and a 30 μA current was passed through each recording site for 2 s to generate small electrolytic lesions (Ugo Basile). Mice were then perfused with 0.9% NaCl. The brains were removed and stored at 4°C, then fixed in 4% PFA for 2 days, rinsed in PBS for 1 day, and cryoprotected in 30% sucrose for at least 2 days. Hemibrains were cut into 30 μm coronal sections with a microtome (Leica) and stored in cryoprotectant at −20°C. Every third right hemibrain section was stained with cresyl violet, then electrolytic lesion locations were observed under a light microscope (Leica). Mice were excluded from electrophysiological analysis if they did not contain at least 2 electrode sites in CA1 pyramidal layer and from CA3 SG power analyses if they had no electrode sites in CA3 (Table S1).

#### Analysis of neural data

Neural data was analyzed with custom software written in MATLAB (Mathworks) with the Chronux toolbox (http://www.chronux.org) and Trodes to MATLAB software (SpikeGadgets). The anatomical location of each electrode site was determined by examining Nissl-stained histological sections, raw LFP traces, the SWR-triggered spectrogram signature, and dentate spikes. Data were band-pass Butterworth filtered at 0.1–300 Hz, then downsampled to 1000 Hz and analyzed as LFP. Raw LFP data were band-pass equiripple filtered at 150–250Hz for SWRs and at 30–50Hz for SG. SWRs were detected on the CA1 site closest to the center of the pyramidal layer and defined by the envelope of the ripple-filtered trace exceeding 5 SD above baseline for at least 15 ms (Cheng and Frank, 2008). Analysis of SWRs was restricted to periods of extended immobility, when the mouse velocity was < 1 cm/s for 30 seconds or more. A recording session was excluded if the mouse was immobile for less than 10 min out of the 60 min session (1/65 and 3/85 sessions for apoE3-KI and apoE4-KI in the screen cohort and 9/65 and 2/80 sessions for apoE3-KI and apoE4-KI 5–8 month recording of the replication cohort). SWR-triggered spectrograms for each electrode site were calculated across all SWRs with the multitaper method, as previously described (Carr et al., 2012), with a 100 ms sliding window. For illustration in figures, a 10 ms sliding window was used (see Figure 1C). SWR-associated SG power was calculated as the averaged z-scored power over the 30–50 Hz frequency band 0–100 ms after ripple detection. This was then averaged over all SWR events and over all electrode sites within that cell layer or subregion. SG power was analyzed for three regions: CA1 stratum radiatum, CA3 including stratum pyramidale and stratum radiatum, and dentate gyrus including hilus and granule cell layers. Only dentate gyrus sites with visually confirmed dentate spikes were included in analysis. All measurements were analyzed per session, then averaged across all sessions to control for any effects of estrous cycling. Thus, each mouse contributed a single number to all comparisons.

#### Analysis of behavioral data

For the screen cohort, we tested, for all hidden days, escape latency, slope of escape latency from day 1 to each day, and difference between escape latency on the last session of a day and the first session of the next day, and for probe days, percent time in target quadrant, number of target crossings, and area under the distance to platform curve. For the MWM task, escape latency during hidden trials and target crossings, percent time in target quadrant, and distance to platform location curve during probe trials were extracted from Noldus (Ethovision). Escape latency was averaged across the 4 daily trials to create 1 value per day. The following two analyses were not provided by Noldus and so were calculated in Excel (Microsoft). Escape latency slope was calculated as the slope of the first-degree polynomial which best fit the average daily escape latencies across a range of hidden days. Area under the distance to platform curve was calculated with the trapezoid method. For the APA task, number of entrances into the shock zone, latency to first entrance to the shock zone, path length relative to the arena, and percent time spent in target opposite the shock zone for each day, plus the number of shocks that would have been received per entrance if the shock were active during the probe trial (pseudoshocks/entrance), were extracted from Tracker software (BioSignal Group). Distance travelled relative to the shock zone per movement bout was extracted using custom Python scripts that analyzed video tracking data extracted from Tracker software (BioSignal Group). When data was non-parametric, they were aligned rank transformed using ARTool (University of Washington).

### Quantification and Statistical Analysis

Performance scores were calculated by taking the z-score of each behavioral measure, inverting the sign of all behavioral measures where higher values indicate worse performance, and averaging across all behavioral measures. For the learning performance score for MWM, all behavioral measures that were significantly correlated with SWR abundance in the screen cohort were used: slope of the escape latency curve across days 1–2 and 1–3, escape latency on day 3, and overnight change in escape latency over days 1–2. For the learning performance score for APA, all behavioral measures that were significantly correlated with SWR abundance in the replication cohort were used: latency to first entrance, path length, and percent time in quadrant opposite the shock zone on day 2. For the memory precision performance score for MWM, all behavioral measures that were significantly correlated with CA3 SG power during SWRs in the screen cohort were used: percent time in target quadrant on probes 1 and 2, number of target crossings on probe 1, and area under the distance to platform curve on probes 1 and 2. For the memory precision performance score for APA, all behavioral measures that were significantly correlated with CA3 SG power during SWRs in the replication cohort were used: number of entrances into the shock zone on day 2 and distance travelled relative to shock zone per movement bout on days 2 and 3.

Statistics were computed using Prism software (Graphpad). Statistical test used, exact n, test statistic values, degrees of freedom, and exact p value are in figure legends. When a central value is plotted, it is always mean ± SEM, as indicated in figure legends. In all cases, n represents number of animals. Significance was established as p < 0.05. No data were excluded based on statistical tests. Subjects were not randomized or stratified. Sample sizes were based on previous studies (Andrews-Zwilling et al., 2010; Gillespie et al., 2016). Most data were normally distributed as shown by Shapiro-Wilk test, and variances between groups were similar as shown by F test. In these instances, we used two-tailed paired and unpaired t tests, two-way ANOVA, two-way mixed-effects analysis (for data sets with missing values), and Pearson correlations. Post-hoc testing was done with Sidak correction for multiple comparisons. When these assumptions were violated, we used two-tailed Mann-Whitney U tests, two-way ANOVA or mixed-effects analysis on aligned rank transformed data (Wobbrock et al., 2011), and Spearman correlations. All correlations were confirmed to be not driven by a single data point by measuring the significance of each relationship after removing each data point in a custom MATLAB script (Mathworks) and confirming p < 0.05.

---

**Figure S1.**
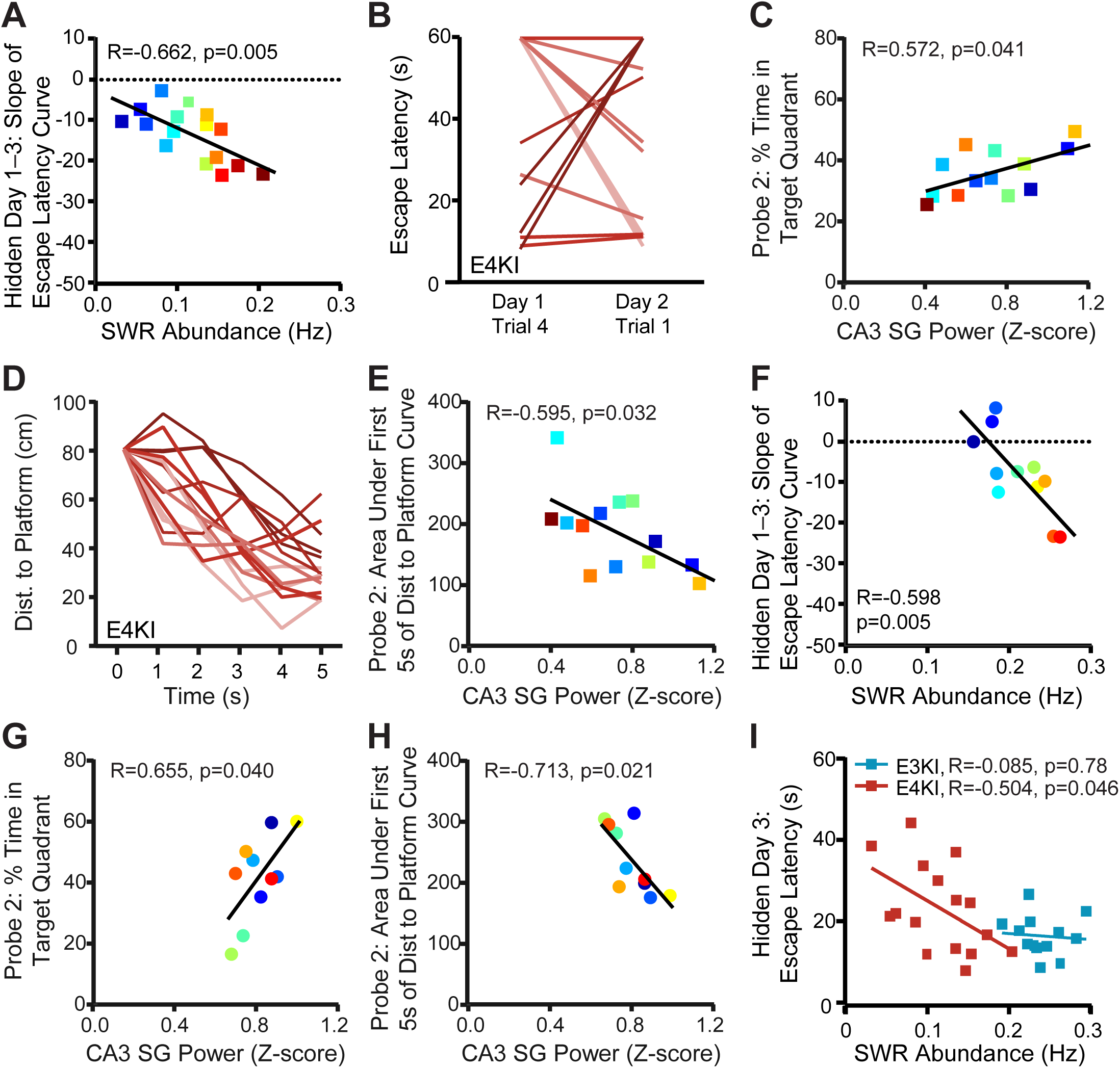
Examples of behavioral metrics used in correlations. Related to Figure 2. (A) SWR abundance predicts slope of escape latency over hidden days 1–3 (F(1,14) = 10.91), n = 16 mice. (B) Individual apoE4-KI mouse escape latency curves over hidden sessions, showing difference between day 1, trial 4 and day 2, trial 1 (overnight), colored from most negative (light) to most positive (dark), n = 16 mice. (C) CA3 SG power during SWRs predicts percent time spent in quadrant that previously contained the platform on probe 2 (F(1,11) = 5.34), n = 13 mice. (D) Individual apoE4-KI mouse distance to platform curves during the first 5 seconds of probe 1, colored from best (light) to worst (dark) cumulative distance to platform over the curve, n = 16 mice. (E) CA3 SG power during SWRs predicts area under the curve of the distance to the prior platform location during the first 5 seconds of probe 2 (F(1,11) = 6.01), n = 13 mice. In A–E, apoE4-KI mice aged 12–18 months at electrophysiological recording and 13–19 months at MWM. (F) In a replication experiment in a separate cohort of animals, SWR abundance predicts slope of escape latency over hidden days 1–3 (F(1,9) = 13.39), n = 11 mice. (G,H) In a replication experiment in a separate cohort of animals, CA3 SG power during SWRs predicts (G) percent time spent in quadrant that previously contained the platform (F(1,8) = 6.02) and (H) area under the curve of the distance to the prior platform location during the first 5 seconds of probe 2 (F(1,8) = 8.27), n = 10 mice. In F–H, apoE4-KI mice aged 13–17 months at electrophysiological recording and 14–18 months at MWM. (I) Example of how SWR abundance for apoE3-KI mice does not overlap with that of apoE4-KI mice and does not predict escape latency for apoE3-KI mice, demonstrating a ceiling effect (F(1,11) = 0.80 for apoE3-KI, n = 13; F(1,14) = 4.77 for apoE4-KI, n = 16). Mice aged 12–18 months at electrophysiological recording and 13–19 months at MWM. Points colored in order of SWR abundance from blue (lowest) to red (highest). Pearson correlations.

**Figure S2.**
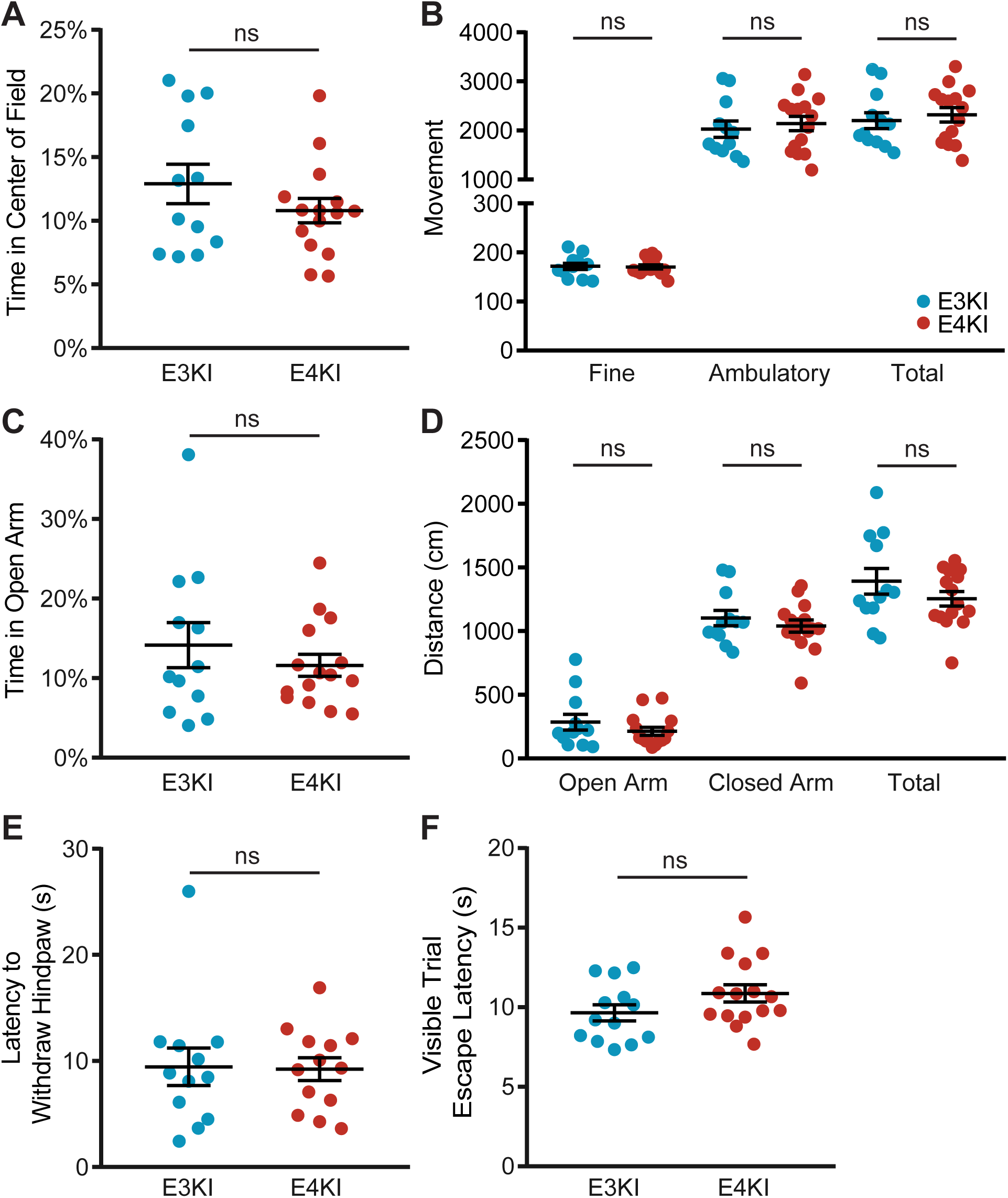
ApoE4-KI mice are not impaired in non-spatial behaviors. Related to Figure 2 and 3. (A) Percent time in center of open field (t(25) = 1.19, p = 0.24). (B) Number of instances of detected movement in the open field (t(25) = 0.19, p = 0.80 for fine; t(25) = 0.52, p = 0.61 for ambulatory; and t(25) = 0.54, p = 0.60 for total). (C) Percent time in open arm of elevated plus maze (t(25) = 0.84, p = 0.41). (D) Distance travelled in elevated plus arms (Mann-Whitney U = 72.5, p = 0.41 for open arms; t(25) = 0.81, p = 0.43 for closed arms; t(25) = 1.25, p = 0.22 for total movement). (E) Latency to withdraw hind paw on hot plate (Mann-Whitney U = 66, p = 0.54). N = 12 apoE3- KI and n = 13 apoE4-KI mice, aged 15–20 months. (F) Escape latency on MWM trials with platform labeled with flag (t(25) = 1.26, p = 0.22). N = 12 apoE3-KI and n = 15 apoE4-KI mice, aged 14–18 months, unless otherwise specified. All tests are unpaired t tests unless otherwise specified. Error bars indicate mean ± SEM.

**Figure S3.**
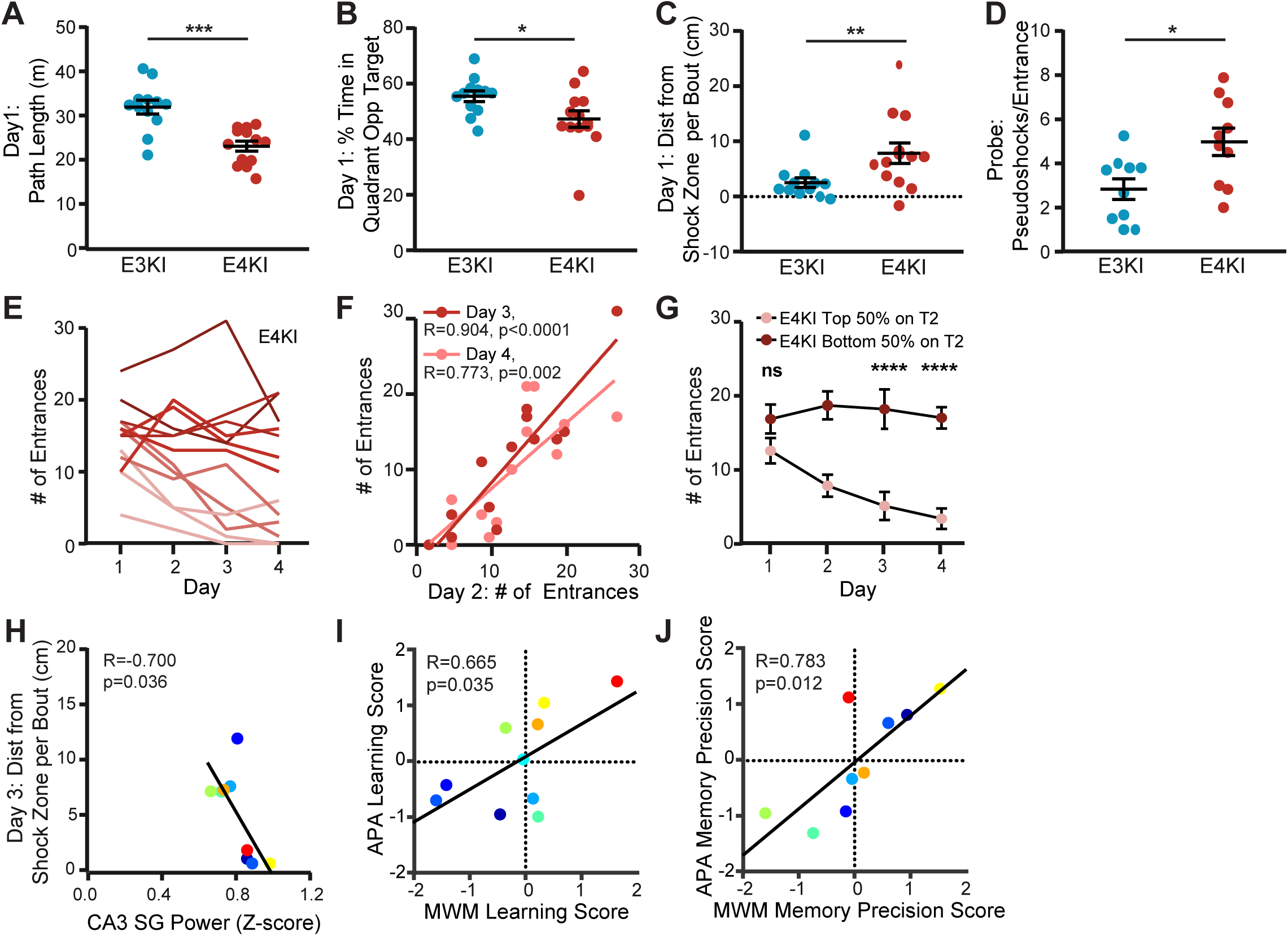
Aged apoE4-KI mice show impaired acquisition of a spatial avoidance task. Related to Figure 3. (A) Total path length on day 1 (unpaired t test, t(23) = 4.64, p = 0.0004). (B) Percent time spent in quadrant opposite the shock zone on day 1 (unpaired t test, t(23) = 2.27, p = 0.033). (C) Distance travelled during each movement bout relative to the shock zone boundary on day 1 (Mann Whitney U = 30, p = 0.008). (D) Shocks that would have been received per shock zone entrance during probe were the field electrified; mice which did not enter the shock zone during probe are excluded (unpaired t test, t(18) = 2.76, p = 0.013). In A–D, n = 12 apoE3-KI mice and n = 13 apoE4-KI mice, aged 15–20 months. (E) ApoE4-KI mice show wide variation in performance. Entrances into the shock zone curves colored from best (light) to worst (dark) average performance over all days. (F) Number of entrances on day 2 predicts number of entrances on days 3 (F(1,11) = 49.27) and 4 (F(1,11) = 16.32). (G) Number of entrances into the shock zone for apoE4-KI mice divided into 2 groups based on number of entrances on day 2. Only differences on days 1, 3, and 4 were examined, yielding no difference in day 1 (t(44) = 1.66, p = 0.36) and significant differences on days 3 (t(44) = 6.06, p < 0.0001) and 4 (t(44) = 5.28, p < 0.0001); n = 7 top 50%, n = 6 bottom 50%. Unpaired t tests with Sidak’s multiple comparison adjustment. (H) CA3 SG power during SWRs predicts distance travelled per movement bout relative to shock zone boundaries on day 3 (F(1,7) = 6.74); n = 9 mice. (I) MWM learning performance score predicts APA learning performance score (F(1,8) = 6.34); n = 10 mice. (J) MWM memory precision performance score predicts APA memory precision performance score (F(1,7) = 11.1); n = 9 mice. In E–J, apoE4-KI mice, aged 15–20 months. All correlations are Pearson correlations. *p < 0.05, **p < 0.01, ***p < 0.001, ****p < 0.0001. Error bars indicate mean ± SEM.

**Figure S4.**
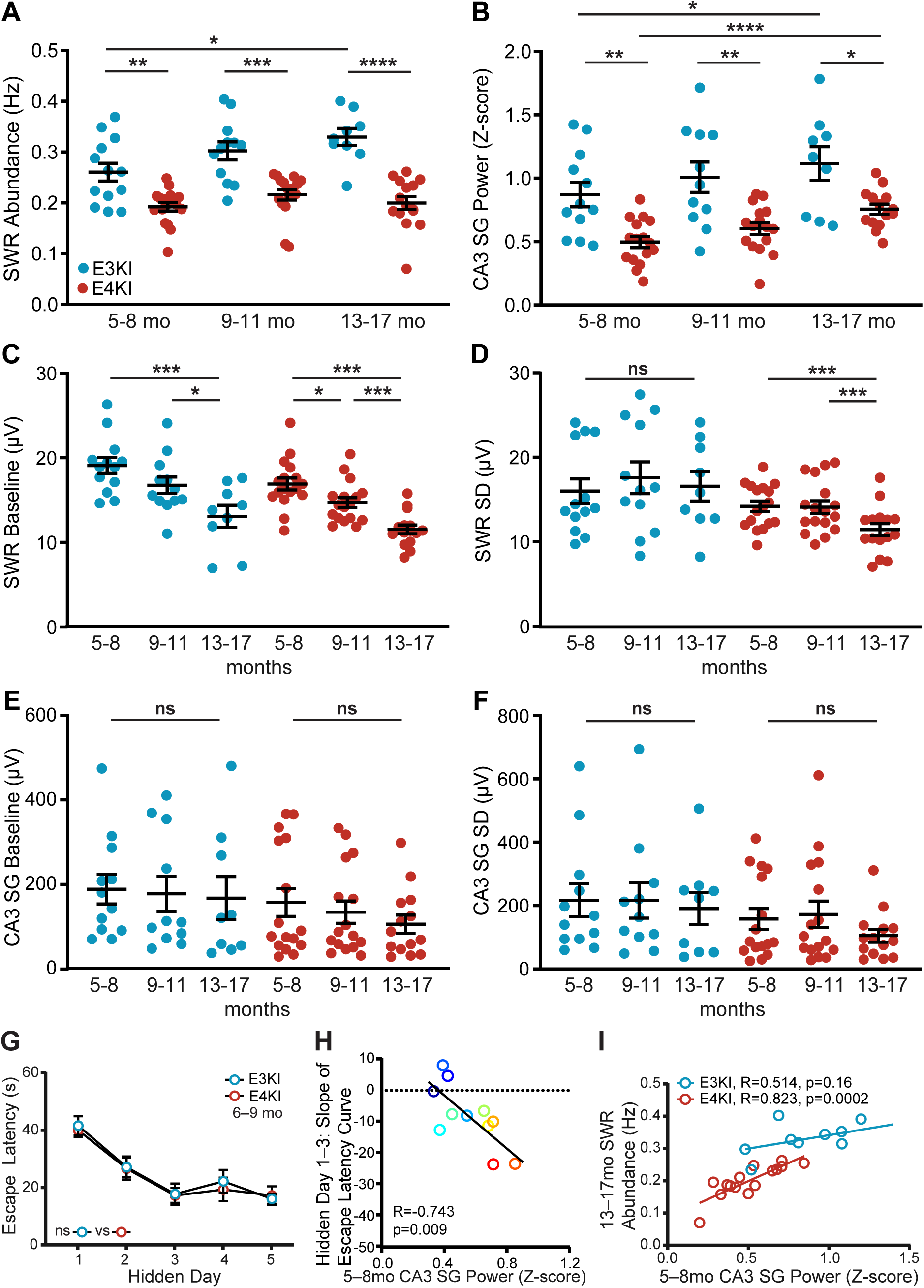
Properties of SWRs and associated SG power in CA3 over aging in apoE3-KI and apoE4-KI mice. Related to Figure 4. (A) SWR abundance of apoE3-KI mice is higher than in apoE4-KI mice across all ages. Two-way mixed-effects analysis of aligned rank transformed data shows significant effect of genotype (F(1,28) = 43.99, p < 0.0001) and post-hoc Mann-Whitney U test with Sidak adjustment shows significant difference at 5–8 months (U = 43, p = 0.012, n = 13 apoE3-KI and n = 17 apoE4-KI mice), 9–11 months (U = 26, p = 0.0012, n = 12 apoE3-KI and n = 17 apoE4-KI mice) and 13–17 months (U = 5, p < 0.0001, n = 9 apoE3-KI and n = 15 apoE4-KI mice). SWR abundance increases over aging in apoE3-KI mice (t(19) = 2.926, p = 0.028, n = 13 for 5–8 months, n = 9 for 13–17 months). (B) CA3 SG power during SWRs in apoE3-KI mice is higher than in apoE4-KI mice across all ages. Two-way mixed-effects analysis of aligned rank transformed data shows significant effect of genotype (F(1,26) = 12.86, p = 0.0014) and post-hoc Mann-Whitney U test with Sidak adjustment shows significant difference at 5–8 months (U = 31, p = 0.0054, n = 12 apoE3-KI and n = 16 apoE4-KI mice), 9–11 months (U = 35, p = 0.024, n = 11 apoE3-KI and n = 16 apoE4-KI mice) and 13–17 months (U = 31, p = 0.0456, n = 9 apoE3-KI and n = 14 apoE4-KI mice). CA3 SG power during SWRs increases over aging in apoE3-KI mice (t(18) = 3.14, p = 0.017, n = 12 for 5–8 months, n = 9 for 13–17 months) and apoE4-KI mice (t(28) = 5.66, p < 0.0001, n = 16 for 5–8 months, n = 14 for 13–17 months). (C) Baseline across the SWR frequency band decreases over aging in apoE3-KI mice (t(19) = 4.60, p = 0.0006, n = 13 for 5–8 months, n = 9 for 13–17 months; t(19) = 2.81, p = 0.033, n = 12 for 9– 11 months, n = 9 for 13–17 months) and apoE4-KI mice (t(30) = 7.25, p < 0.0001, n = 17 for 5–8 months, n = 15 for 13–17 months; t(30) = 3.10, p = 0.012, n = 17 for 5–8 months, n = 7 for 9–11 months; t(30) = 4.26, p = 0.0006, n = 17 for 9–11 months, n = 15 for 13–17 months). (D) SD across the SWR frequency band decreases over aging in apoE4-KI mice (t(30) = 4.34, p = 0.0004, n = 17 for 5–8 months, n = 15 for 13–17 months; t(30) = 4.18, p = 0.0007, n = 17 for 9– 11 months, n = 15 for 13–17 months), but not apoE3-KI mice (t(19) = 0.25, p = 0.99, n = 13 for 5–8 months, n = 9 for 13–17 months). (E) Baseline across SG frequency band in CA3 does not change over aging in apoE3-KI mice (t(18) = 0.18, p = 1.0, n = 12 for 5–8 months, n = 9 for 13–17 months) or apoE4-KI mice (Mann Whitney test with Sidak adjustment, U = 88, p = 0.70, n = 16 for 5–8 months, n = 14 for 13–17 months). (F) SD across SG frequency band in CA3 does not change over aging in apoE3-KI mice (t(18) = 0.50, p = 0.95, n = 12 for 5–8 months, n = 9 for 13–17 months) or apoE4-KI mice (Mann Whitney U test with Sidak adjustment, U = 98, p = 0.93, n = 16 for 5–8 months, n = 14 for 13–17 months). In A–F, all comparisons unpaired t tests with Sidak adjustment unless otherwise specified. (G) ApoE4-KI mice show no impairment on MWM at ages 6–9 months, n = 13 apoE3-KI and n = 13 apoE4-KI mice. (H) CA3 SG power during SWRs measured at 5–8 months predicts slope of escape latency over hidden days 1–3 (F(1,9) = 10.89 on MWM task at 14–18 months, n = 11 apoE4-KI mice, Pearson correlation. (I) CA3 SG power during SWRs measured at 5–8 months predicts SWR abundance in the same mouse at 13–17 months for apoE4-KI (F(1,13) = 27.19, n = 15) but not apoE3-KI mice (F(1,7) = 2.52, n = 9), Pearson correlations. *p < 0.05; **p < 0.01; ***p < 0.001; ****p < 0.0001. Error bars indicate mean ± SEM.

**Table S1.** Sample sizes for all experiments. Related to Methods.

| Age | Metric | Cohort 1 |  | Cohort 2 |  |
| --- | --- | --- | --- | --- | --- |
|  |  | apoE3-KI | apoE4-KI | apoE3-KI | apoE4-KI |
| 5–8 months | SWR Abundance | -- | -- | 13 | 17<sup>a</sup> |
|  | CA3 SG Power | -- | -- | 12 | 16 |
|  | MWM | -- | -- | 13 | 13 |
| 9–11 months | SWR Abundance | -- | -- | 12<sup>b</sup> | 17 |
|  | CA3 SG Power | -- | -- | 11<sup>b</sup> | 16 |
| 12–20 months | SWR Abundance | 13 | 16 | 9<sup>c</sup> | 15<sup>d</sup> |
|  | CA3 SG Power | 11 | 13 | 9<sup>c</sup> | 14<sup>d</sup> |
|  | MWM, Open Field, Elevated Plus | 20 | 19 | 12<sup>e</sup> | 15<sup>f</sup> |
|  | APA, Hot Plate | -- | -- | 12<sup>e</sup> | 13<sup>g</sup> |

<sup>a</sup>4 apoE4-KI mice aged 7 months were added to the study after the first MWM had been completed.
<sup>b</sup>Implant no longer functional in 1 apoE3-KI mouse
<sup>c</sup>Implant no longer functional in 2 apoE3-KI mice, 1 apoE3-KI mouse died
<sup>d</sup>Implant no longer functional in 1 apoE4-KI mouse, 1 apoE4-KI mouse died
<sup>e</sup>3 apoE3-KI mice died, 3 apoE3-KI mice added
<sup>f</sup>4 apoE4-KI mice died, 3 apoE4-KI mice added
<sup>g</sup>1 apoE4-KI mouse (added just for behavior) excluded due to motor deficits, 1 apoE4-KI mouse died

**Table S2.** SWR abundance and associated CA3 SG power correlations with MWM metrics identified in screen cohort. Related to Figure 2.

| Metric | SWR Abundance |  |  |  | CA3 SG Power |  |  |  |
| --- | --- | --- | --- | --- | --- | --- | --- | --- |
|  | R | F | DFn,<br>DFd | P<br>value | R | F | DFn,<br>DFd | P<br>value |
| Slope of Escape Latency Days 1-2 | **-0.56** | **6.41** | **1, 14** | **0.024** | 0.28 | 0.94 | 1, 11 | 0.35 |
| Slope of Escape Latency Days 1-3 | **-0.66** | **10.91** | **1, 14** | **0.005** | 0.48 | 3.29 | 1, 11 | 0.10 |
| Escape Latency Day 3 | **-0.50** | **4.77** | **1, 14** | **0.046** | 0.16 | 0.27 | 1, 11 | 0.61 |
| Night 1 Learning | **-0.64** | **9.76** | **1, 14** | **0.008** | 0.38 | 1.84 | 1, 11 | 0.20 |
| Probe 1 % in Target Quadrant | -0.11 | 0.16 | 1, 14 | 0.69 | **0.77** | **15.75** | **1, 11** | **0.002** |
| Probe 2 % in Target Quadrant | -0.03 | 0.02 | 1, 14 | 0.90 | **0.57** | **5.34** | **1, 11** | **0.041** |
| Probe 1 Target Crossings | 0.12 | 0.21 | 1, 14 | 0.65 | **0.78** | **17.30** | **1, 11** | **0.002** |
| Probe 1 Dist to Platform Curve | 0.10 | 0.15 | 1, 14 | 0.71 | **-0.61** | **6.55** | **1, 11** | **0.027** |
| Probe 2 Dist to Platform Curve | 0.03 | 0.02 | 1, 14 | 0.90 | **-0.60** | **6.01** | **1, 11** | **0.032** |

Bold values indicate significant correlations.

**Table S3.** All APA metric correlations tested in replication cohort. Related to Figure 3.

| Metric | Trial | SWR Abundance |  |  |  | CA3 SG Power |  |  |  |
| --- | --- | --- | --- | --- | --- | --- | --- | --- | --- |
|  |  | R | F | DFn,<br>DFd | P<br>value | R | F | DFn,<br>DFd | P<br>value |
| # of Entrances | 1 | -0.27 | 0.64 | 1, 8 | 0.45 | -0.24 | 0.43 | 1, 7 | 0.53 |
|  | 2 | -0.09 | 0.07 | 1, 8 | 0.80 | **-0.91** | **32.5** | **1, 7** | **0.0007** |
|  | 3 | -0.14 | 0.16 | 1, 8 | 0.70 | **-0.76** | **9.49** | **1, 7** | **0.02** |
|  | 4 | 0.19 | 0.29 | 1, 8 | 0.61 | **-0.72** | **7.48** | **1, 7** | **0.03** |
| Latency to 1st Entrance | 1 | -0.43 | 1.78 | 1, 8 | 0.22 | 0.01 | 0 | 1, 7 | 0.98 |
|  | 2 | **0.65** | **5.81** | **1, 8** | **0.043** | 0.27 | 0.53 | 1, 7 | 0.49 |
|  | 3 | 0.16 | 0.22 | 1, 8 | 0.65 | 0.10 | 0.08 | 1, 7 | 0.79 |
|  | 4 | 0.36 | 1.19 | 1, 8 | 0.31 | 0.37 | 1.12 | 1, 7 | 0.33 |
| Path Length | 1 | 0.14 | 0.15 | 1, 8 | 0.71 | 0.00 | 0 | 1, 7 | 1 |
|  | 2 | **0.82** | **16.42** | **1, 8** | **0.004** | -0.03 | 0.01 | 1, 7 | 0.93 |
|  | 3 | 0.23 | 0.44 | 1, 8 | 0.52 | -0.28 | 0.61 | 1, 7 | 0.46 |
|  | 4 | 0.53 | 3.08 | 1, 8 | 0.12 | -0.25 | 0.45 | 1, 7 | 0.53 |
| % Time in Opposite Quadrant | 1 | 0.22 | 0.39 | 1, 8 | 0.55 | 0.45 | 1.77 | 1, 7 | 0.22 |
|  | 2 | **0.69** | **7.33** | **1, 8** | **0.027** | 0.22 | 0.37 | 1, 7 | 0.56 |
|  | 3 | -0.10 | 0.07 | 1, 8 | 0.79 | 0.13 | 0.12 | 1, 7 | 0.74 |
|  | 4 | -0.03 | 0.01 | 1, 8 | 0.93 | 0.07 | 0.04 | 1, 7 | 0.85 |
| Dist from Shock Zone per Bout | 1 | -0.30 | 0.77 | 1, 8 | 0.41 | -0.17 | 0.21 | 1, 7 | 0.66 |
|  | 2 | -0.08 | 0.05 | 1, 8 | 0.83 | **-0.75** | **9.04** | **1, 7** | **0.02** |
|  | 3 | 0.05 | 0.02 | 1, 8 | 0.90 | **-0.7** | **6.74** | **1, 7** | **0.04** |
|  | 4 | 0.38 | 1.31 | 1, 8 | 0.29 | -0.15 | 0.15 | 1, 7 | 0.71 |

Bold values indicate significant correlations.

**Table S4.** All behavioral correlations tested with 5–8 month apoE4-KI mouse electrophysiological data. Related to Figure 4.

| Age and Behavior | Metric | SWR Abundance |  |  |  | CA3 SG Power |  |  |  |
| --- | --- | --- | --- | --- | --- | --- | --- | --- | --- |
|  |  | R | F | DFn, DFd | P value | R | F | DFn, DFd | P value |
| 6-9 mo MWM | Slope of Escape Latency Days 1-2 | 0.05 | 0.02 | 1,11 | 0.88 | 0.06 | 0.02 | 1,10 | 0.85 |
|  | Slope of Escape Latency Days 1-3 | 0.17 | 0.33 | 1, 11 | 0.57 | -0.04 | 0.02 | 1, 10 | 0.90 |
|  | Escape Latency Day 3 | -0.03 | 0.01 | 1, 11 | 0.93 | 0.04 | 0.02 | 1, 10 | 0.90 |
|  | Night 1 Learning | 0.22 | 0.55 | 1, 11 | 0.47 | -0.01 | 0.001 | 1, 10 | 0.97 |
|  | Probe 1 % in Target Quadrant | -0.23 | 0.61 | 1, 11 | 0.45 | -0.05 | 0.02 | 1, 10 | 0.88 |
|  | Probe 2 % in Target Quadrant | -0.06 | 0.04 | 1, 11 | 0.84 | 0.09 | 0.08 | 1, 10 | 0.79 |
|  | Probe 1 Target Crossings | -0.48 | 3.36 | 1, 11 | 0.09 | -0.23 | 0.56 | 1, 10 | 0.47 |
|  | Probe 1 Dist to Platform Curve | 0.27 | 0.88 | 1, 11 | 0.37 | 0.14 | 0.19 | 1, 10 | 0.68 |
|  | Probe 2 Dist to Platform Curve | -0.31 | 1.18 | 1, 11 | 0.30 | -0.21 | 0.44 | 1, 10 | 0.52 |
| 14-18 mo MWM | Slope of Escape Latency Days 1-2 | 0.20 | 0.42 | 1,10 | 0.51 | **-0.63** | **5.82** | **1,9** | **0.039** |
|  | Slope of Escape Latency Days 1-3 | -0.11 | 0.12 | 1, 10 | 0.74 | **-0.74** | **10.89** | **1, 9** | **0.009** |
|  | Escape Latency Day 3 | 0.19 | 0.37 | 1, 10 | 0.56 | **-0.74** | **11.08** | **1, 9** | **0.009** |
|  | Night 1 Learning | -0.16 | 0.27 | 1, 10 | 0.62 | -0.45 | 2.30 | 1, 9 | 0.16 |
|  | Probe 1 % in Target Quadrant | -0.49 | 3.20 | 1, 10 | 0.10 | -0.35 | 1.28 | 1, 9 | 0.29 |
|  | Probe 2 % in Target Quadrant | -0.31 | 1.08 | 1, 10 | 0.32 | -0.07 | 0.05 | 1, 9 | 0.83 |
|  | Probe 1 Target Crossings | -0.43 | 2.21 | 1, 10 | 0.17 | -0.28 | 0.77 | 1, 9 | 0.40 |
|  | Probe 1 Dist to Platform Curve | 0.45 | 2.53 | 1, 10 | 0.14 | 0.22 | 0.45 | 1, 9 | 0.52 |
|  | Probe 2 Dist to Platform Curve | 0.17 | 0.29 | 1, 10 | 0.60 | 0.37 | 1.40 | 1, 9 | 0.27 |
| 15-20 mo APA | # of Entrances | 0.60 | 5.09 | 1, 9 | 0.05 | -0.09 | 0.07 | 1, 8 | 0.80 |
|  | Latency to 1<sup>st</sup> | -0.29 | 0.81 | 1, 9 | 0.39 | **0.79** | **13.21** | **1, 8** | **0.007** |
| Trial 2 | Entrance |  |  |  |  |  |  |  |  |
|  | Path Length | 0.01 | 0.0003 | 1, 9 | 0.99 | **0.68** | **6.74** | **1, 8** | **0.03** |
|  | % Time in Opposite Quadrant | -0.18 | 0.31 | 1, 9 | 0.59 | 0.59 | 4.25 | 1, 8 | 0.07 |
|  | Dist from Shock Zone per Bout | 0.46 | 2.48 | 1,9 | 0.15 | 0.11 | 0.09 | 1,8 | 0.77 |
| Trial 3 | Dist from Shock Zone per Bout | 0.43 | 2.02 | 1,9 | 0.19 | 0.20 | 0.33 | 1,8 | 0.58 |

Bold values indicate significant correlations.

